# *Fosl2* regulates the transition from parietal epithelial cells to myofibroblasts in the kidney

**DOI:** 10.1101/2025.11.10.687667

**Authors:** Robert Bronstein, Monica P. Revelo, Fatima Sheikh, John D. Haley, Vivette D’Agati, David J. Salant, John C. He, Yiqing Guo, Sandeep K. Mallipattu

**Author notes:** To whom correspondence should be addressed: Sandeep K. Mallipattu and Robert Bronstein, Department of Medicine/Nephrology, Stony Brook University, 100 Nicolls Road, HSCT16-080E, Stony Brook, NY, USA.

## Abstract

Activation and proliferation of parietal epithelial cells (PECs), located along the inner rim of Bowman’s capsule, drives disease progression in subtypes of glomerulonephritis and focal segmental glomerulosclerosis. In examining the mechanisms contributing to PEC activation two established mouse models were utilized in this study, nephrotoxic serum nephritis (transient model) and podocyte-specific *Klf4* knockout (progressive model). A role for transcription factor FRA2 (*Fosl2*) was uncovered through single nuclear multiomic approaches relating to the regulation of PEC transcriptional/chromatin dynamics. Co-immunoprecipitation followed by mass spectrometry assessed the FRA2 protein interactome in cultured PECs, revealing a potential role for FRA2 in alternative splicing. *Fosl2* expression was then blunted through CRISPR-Cas9 gene editing in cultured PECs, revealing reduced proliferative capacity and the downregulation of myofibroblast markers. *In-vivo* genetic lineage tracing of PECs after nephrotoxic serum revealed PEC-to-myofibroblast trans-differentiation events. Finally, immunostaining of human kidney biopsies with varied subtypes of glomerulonephritis confirmed *Fosl2* expression in activated PECs within crescentic lesions, with single cell deconvolution strategies assigning PEC-skewed proportion ratios to bulk RNA-seq patient data from the NEPTUNE consortium. These results suggest that FRA2 (*Fosl2*) directs a conserved molecular program of PEC-specific responses in subtypes of glomerulonephritis and focal segmental glomerulosclerosis.

## INTRODUCTION

Parietal epithelial cells (PECs) in the kidney glomerulus reside along the inner rim of Bowman’s capsule, performing homeostatic functions under normal physiological conditions, while being the primary constituents of glomerular crescentic and pseudo-crescentic lesions in the context of subtypes of glomerulonephritis and focal segmental glomerulosclerosis (FSGS) (1–5). Given that the aforementioned pathologies are very likely to progress to end-stage kidney disease (ESKD) without treatment, the development of novel therapeutic strategies is critical (4, 6, 7). Furthermore, a comprehensive understanding of PEC activation dynamics across these diseases is necessary to identify novel targets for therapy. Though several potential mediators might trigger the activation of PECs due to podocyte injury and loss (8), it remains unclear how this contributes to potential repair as compared to subsequent PEC proliferation and progression to glomerulosclerosis. While podocyte loss is the inciting event in subtypes of glomerulonephritis and FSGS, subsequent PEC activation and proliferation remains a universal feature critical to the development of eventual glomerulosclerosis (3, 9–11). In this study, we utilized two murine models of rapid podocyte loss and PEC activation and proliferation, nephrotoxic serum nephritis (12–14) (transient model) and mice with podocyte-specific loss of *Krüppel-like factor 4 (Klf4*^Δ^*^Pod^)* (14, 15) (progressive model) to examine shared signaling pathways, gene expression, and chromatin dynamics across activated PEC subgroups that contribute to the progression of disease. Specifically, a combination of single nucleus RNA sequencing (snRNA-seq) and <u>A</u>ssay for <u>T</u>ransposase-<u>A</u>ccessible <u>C</u>hromatin with sequencing (snATAC-seq) were initially conducted across these models to identify unbiased key inter- and intra-cellular signaling pathways as well as putative gene/chromatin regulatory factors critical for the activation of PECs and subsequent sclerosis. We identified the transcription factor (TF) FRA2 (gene name: FOS Like Antigen 2 – *Fosl2*), whose gene expression and genomic distribution of consensus binding motifs were significantly enriched in activated PEC cohorts. Additionally, we identified a potentially novel autocrine/paracrine ligand (*Secreted phosphoprotein 1, Spp1*) – receptor (*Cd44*) interaction (16–19) within the activated PEC cluster, which acts as a positive feedforward driver of PEC activation, proliferation and glomerulosclerosis. Rapid Immunoprecipitation Mass Spectrometry of Endogenous Proteins (RIME) (20) was conducted in cultured mouse PECs to show the FRA2-centric protein interactome – wherein we determined that FRA2 binds to numerous RNA-binding proteins (RBPs) and other cogs of the mRNA alternative splicing machinery. Interestingly, analysis of changes in transcriptome between the PEC and myofibroblast clusters led us to postulate that PECs may serve as reservoirs for eventual myofibroblast seeding of the glomerular lesion space, which was corroborated through *in-vivo* lineage tracing. Finally, we utilized single-cell deconvoluted bulk RNA-seq data from human kidney biopsies with FSGS, demonstrating the presence of molecularly similar activated PEC/myofibroblast populations, with subsequent immunofluorescence of human kidney biopsy samples with subtypes of FSGS and glomerulonephritis, confirming FRA2 (*Fosl2*) expression in activated PECs within crescentic lesions.

## RESULTS

### Mouse kidney Multiomics in the nephrotoxic serum nephritis murine model identified key intercellular signaling pathways in activated PECs

To identify key intercellular signaling pathways in activated PECs, we initially utilized mice at day 7 and 14 post-NTS treatment (**Figure 1A**). As previously reported in the nephrotoxic serum (NTS) nephritis model (15), albuminuria was significantly increased at day 7 and 14 post-NTS as compared to IgG control but was reduced from day 7 to 14 post-NTS treatment (**Figure 1A**). Periodic acid-Schiff (PAS) staining shows a significant increase in the percentage of crescents, FSGS lesions, tubular injury, and interstitial inflammation in both day 7 and 14, which were reduced at day 14 as compared to day 7 post-NTS treatment (**Figure 1A & Supplemental Table 1**). To investigate concurrent changes to transcriptome and chromatin accessibility in PECs specifically, we conducted 10X Multiomics (snRNA & ATAC-seq) at day 7 and 14 post-NTS as compared to IgG control (**Supplemental Figure 1**). Cluster Uniform Manifold Approximations and Projections (UMAPs) of the aggregate time course dataset revealed the presence of an emergent activated PEC cluster displaying differential gene expression and chromatin accessibility enrichment, which included canonical PEC activation marker, *Cd44*, and the mesenchymal marker, *Vcam1* (**Figures 1B,C & Supplemental Figure 2B**). The top 10 differentially enriched genes (DEGs) in the PEC cluster revealed the well-known ligand-receptor (L/R) duo of *Spp1*/*Cd44* (21), in addition to genes such as *Dock10* (**Supplemental Figure 2C, Supplemental Tables 2 and 3**), which is a guanine nucleotide exchange factor (GEF) for the Rho GTPase family of proteins, whose critical signaling functions impinge upon intracellular cytoskeletal dynamics (22, 23) (**Figure 1C, Supplemental Figure 2C, Supplemental Tables 2 and 3**). Additionally, *Havcr1* (KIM-1) was found to be enriched in activated PECs, which is a well described kidney injury marker predominantly reported in PT and possibly indicating the presence of dedifferentiating PEC sub-populations (**Supplemental Figure 2A**) (24). Gene Set Enrichment Analysis (GSEA) of enriched PEC-cluster genes with the WikiPathways software package (25) revealed strong concordance of upregulated PEC genes with canonical Primary FSGS gene signatures. The downregulated PEC genes were involved in metabolic cellular signaling cascades such as that of amino acid metabolism and PPAR signaling (**Figure 1D**). REACTOME pathway (25) analysis additionally showed an enrichment of mitotic and RHO GTPase gene categories in the upregulated PEC genes and validated the reduction in metabolic pathways in the downregulated PEC genes (**Figure 1E**). To validate a potential interaction between PEC-enriched *Spp1* and *Cd44* more systematically, we employed the CellChat (26) R package (**Figure 1F & Supplemental Figures 3A-H**). CellChat computed communication probabilities identified a strong, putatively autocrine, L/R loop specific to PECs (**Figure 1F**). While the dominant sender/receiver plots identified the Loop of Henle as having the most overall L/R links across kidney clusters, the *Spp1*/*Cd44* pair was most enriched in the PEC cluster (**Figure 1F**). All other potential *Cd44*-specific ligands were much more prominently invoked via the snRNA-seq data as belonging to non-PEC ligand-receptor loops (**Supplemental Figures 3A-H**). To investigate how the *Spp1*/*Cd44* ligand-receptor loop was temporally constructed in PEC populations *in-vivo*, we quantified the multiome-derived spliced and unspliced mRNA transcript counts and leveraged RNA Velocity analysis (27). This revealed an overwhelming presence of spliced vs unspliced *Spp1* transcripts, suggesting it was likely turned on earlier than *Cd44* in PEC populations (**Figure 1G**). Triple immunofluorescence staining for OPN (*Spp1*), CD44 and SSeCKS (*Akap12* – canonical PEC marker (28, 29)) displayed a robust co-localization of OPN and CD44 in PECs along Bowman’s capsule (**Figure 1H**). Collectively, these data point to a potential L/R signaling loop which may be instrumental in PEC activation, crescent development, and ultimately glomerulosclerosis - cell behaviors that were additionally illuminated by further GSEA GO and KEGG ontology and pathway analyses, respectively (**Supplemental Figures 4A-D**). Genes showing enrichment in the PEC cluster were characterized as having roles in cytoskeletal and cell leading edge behaviors (**Supplemental Figure 4C**), with other gene sets responsible for varying metabolic functions being turned down in the activated PEC population (**Supplemental Figure 4B**).

**Figure 1:**
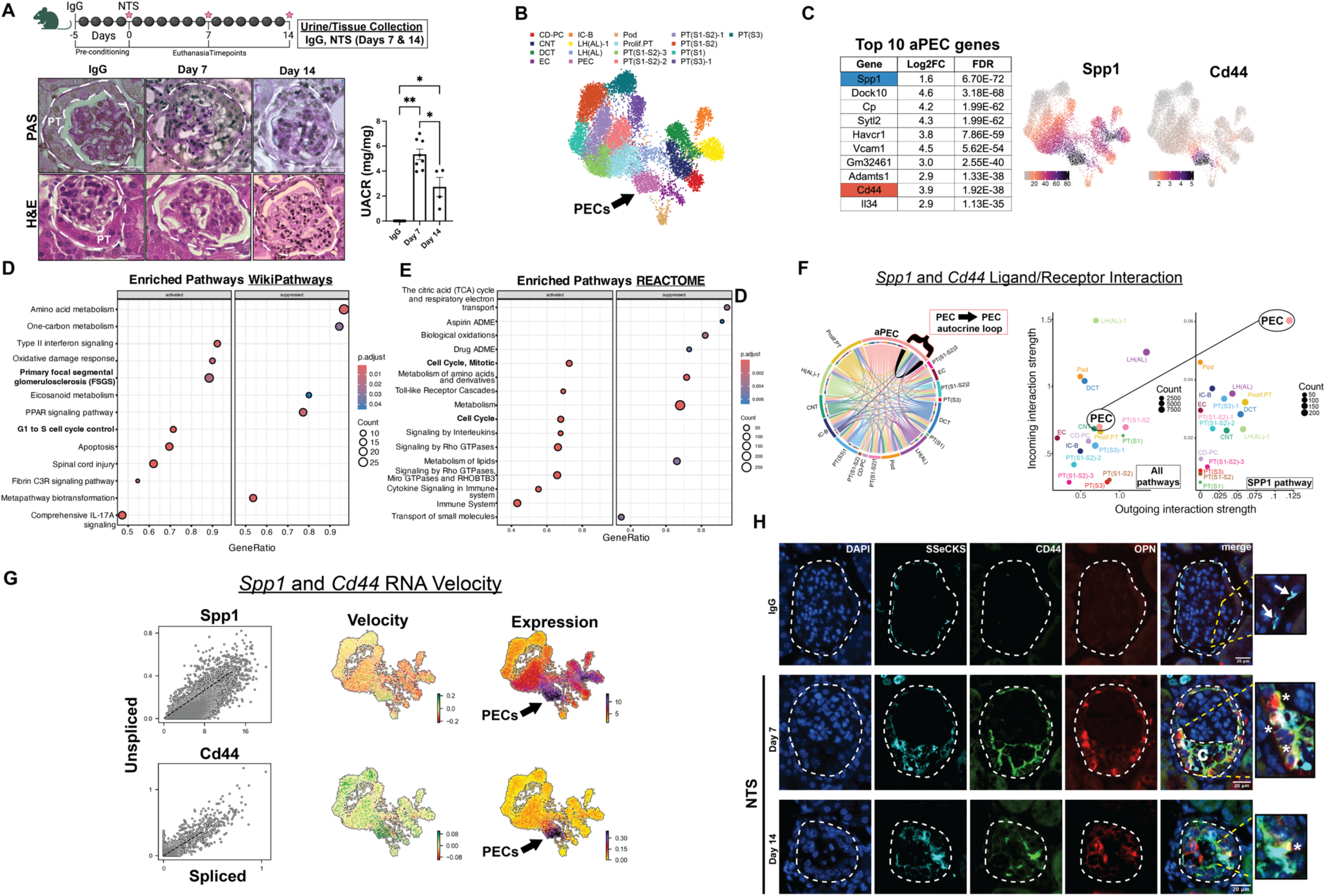
Mouse kidney Multiomics in NTS-induced nephritis identified key intercellular signaling pathways in activated PECs. **(A)** Schematic of nephrotoxic serum (NTS) nephritis model, representative images of PAS staining, and urine albumin to creatinine (UACR) measurements along the experimental time-course. The dotted circles show the glomerular region across all images, with PT in IgG images representing first segment of proximal tubule. **(B)** UMAP of the total NTS dataset (IgG, Day 7, Day 14) totaling 17 distinct cell-type clusters. **<u>Cell-type abbreviations</u>:** PECs, parietal epithelial cells; Pod, podocytes; PT, proximal tubule; Prolif.PT, proliferating proximal tubule; LH(AL), loop of Henle (ascending limb); LH(AL)-1, loop of Henle ascending limb subcluster; DCT, distal convoluted tubule; CNT, connecting tubule; CD-PC, collecting duct principal cells; IC-B, collecting duct intercalated cells (type B); EC, endothelial cells. **(C)** Top 10 PEC-enriched genes obtained by comparing PEC clusters with all other clusters. *Spp1* and *Cd44* UMAP feature plots displaying PEC-specific enrichment of this ligand-receptor pair. **(D)** WikiPathways GSEA ridge plot of top up/down-regulated genes enriched in the PEC cluster**. (E)** REACTOME enriched pathway dot plot of top up/down-regulated genes enriched in the PEC cluster. **(F)** CellChat chord diagram displaying *Spp1*/*Cd44*-specific ligand-receptor loops, including putative PEC-to-PEC autocrine loop. Dominant sender (sources) and receiver (targets) scatter plots: (left) enrichment of ligand-receptor pairs across all clusters and pathways, (right) enrichment of *Spp1*/*Cd44* ligand-receptor pair across all clusters. **(G)** ‘Steady-state’ spliced vs unspliced transcript ratio plots for *Spp1*/*Cd44* ligand-receptor pair along with UMAP embeddings overlayed independently with RNA velocity and gene expression values for both *Spp1* and *Cd44.* **(H)** Representative images of immunofluorescence staining in NTS-treated (day 7 & day 14 with **C** labeling crescentic area) as compared to IgG-treated mice for: SSeCKs (*Akap12*), CD44 & OPN (*Spp1*). Arrows in IgG row point to presence of quiescent SSeCKs+ PECs, with asterisks at days 7 & 14 denoting triple positive cells (SSeCKs+/CD44+/OPN+).

### Nephrotoxic serum-induced nephritis transcriptome implicates emergence of transitional PEC/myofibroblast cell states

Seurat cluster UMAPs derived from QC normalized aggregate samples (**Supplemental Figures 5A-E**) demonstrated that the PEC population is most abundant at day 7 as compared to IgG control and day 14 post-NTS (**Figure 2A**). While *Spp1* is expressed in other clusters, it is mostly revealed as a bifurcated ridge of low/high expressing cells, with all emergent PEC populations showing only high expression (**Figure 2B**). Furthermore, the total number of PECs decreased from day 7 to 14 post-NTS, but the *Spp1* expression remained elevated at day 14 – indicating a potential role for this PEC-secreted ligand in both PEC activation and eventual glomerulosclerosis (**Figure 2C**). Sub-clustering this population resulted in five distinct PEC subgroups (**Figures 2D-E**), with co-enrichment for both *Spp1* and *Cd44* in PEC sub-cluster 3 (**Figure 2F, Supplemental Table 4**). Additionally, *Spp1* and *Cd44* were not only expressed in the same PEC sub-type, but actually co-expressed in the same cells within PEC sub-cluster 3 (**Figure 2G**), further corroborating our CellChat results (**Figure 1D**). WikiPathways and REACTOME pathway analyses of PEC sub-clusters 2/3 DEGs (**Figure 2H)** uncovered a mix of terms related to cell focal adhesion, chemotaxis and primary FSGS gene expression profiles (**Supplemental Figures 6-8**). Furthermore, pseudo-temporal ordering using Monocle3 of the PEC sub-cluster space and the adjoining bifurcation of the kidney proliferative PEC/myofibroblast clusters demonstrated pseudotime trajectories from the *Spp1*/*Cd44* co-expressing cells to both forked cluster endpoints (proliferative PECs and myofibroblasts) (**Figure 2I**), as supported by the lineage paths taken towards the bifurcation from the starting root cells (**Figure 2J**). Monocle3 trajectories from these PEC *Spp1*/*Cd44* co-expressing cells revealed marker genes specific to the two branch points (e.g., *Mki67, Col1a1*) (**Figure 2K & Supplemental Figure 9**). Monocle3 modules 1-5, representing distinct points along the pseudotemporal trajectory, were indicative of the potential relationship between PECs and proliferative PEC/myofibroblasts (**Figures 2L,M**). WikiPathway analysis of modules 2 and 4 recapitulated our previous PEC subcluster analysis (**Figure 2H**), indicating a transition between sub-clusters of PECs to proliferative PECs and myofibroblasts post-NTS (**Figure 2N**).

**Figure 2:**
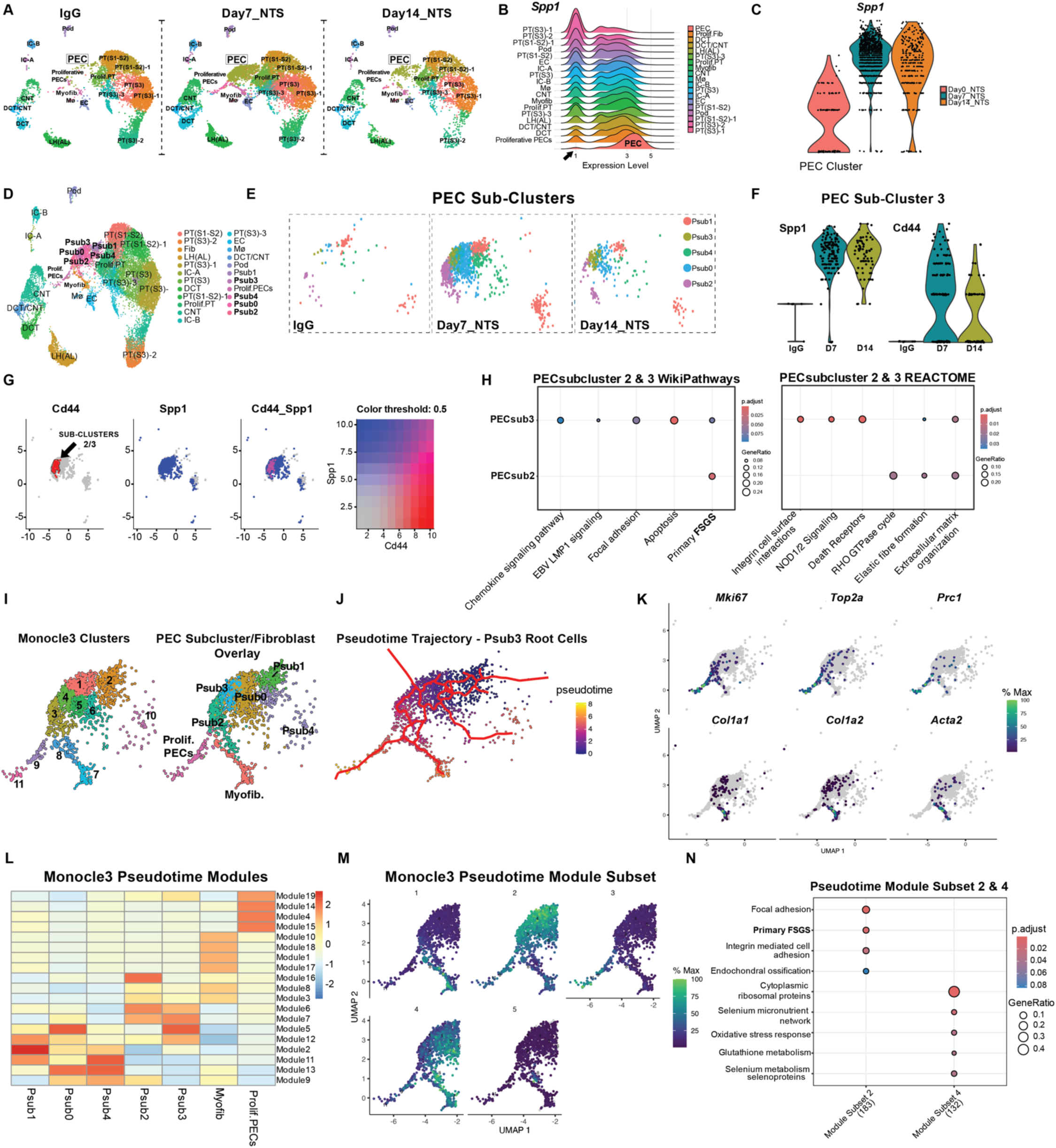
NTS-induced nephritis transcriptome shows emergence of transitional PEC/fibroblast cell states. **(A)** Seurat RNA UMAPs representing IgG, 7 & 14 post-nephrotoxic serum (NTS). **<u>Cell-type</u> <u>abbreviations</u>:** PECs, parietal epithelial cells; Proliferative PECs, cycling parietal epithelial cells; Pod, podocytes; PT, proximal; Prolif.PT, proliferating; LH(AL), loop of Henle (ascending limb); DCT, distal convoluted tubule; CNT, connecting tubule; DCT/CNT, distal convoluted tubule/connecting tubule transitional cells; IC-A, collecting duct intercalated cells (type A); IC-B, collecting duct intercalated cells (type B); EC, endothelial cells; Myofib/Mφ, myofibroblasts/macrophages. **(B)** Ridge plot of *Spp1* gene expression distribution across aggregate NTS time-course clusters. **(C)** Violin plot of *Spp1* expression in the PEC cluster across the three different timepoints in the NTS time-course. **(D)** Aggregate UMAP displaying the relative positioning of emergent PEC sub-clusters, following re-clustering of cells only within the PEC cluster. **(E)** Breakdown of PEC sub-clusters from IgG, 7 & 14 post-NTS, in the absence of all other kidney cluster UMAP embeddings. **(F)** *Spp1* and *Cd44* gene expression violin plots for PEC sub-cluster 3 **(G)** Blended UMAP embedding of *Spp1* (purple) and *Cd44* expression (red) largely overlapping in PEC sub-clusters 2 & 3 (as per the black arrow). **(H)** Pathway analysis results for PEC subclusters 2 & 3 through WikiPathways and REACTOME. **(I)** UMAPs of PEC subclusters with the addition of myofibroblast and proliferative PEC clusters - broken down by Monocle3-called clusters (left) and Seurat-called clusters (right). **(J)** Monocle3 trajectory map of PEC/Myofibroblast/Proliferative PEC cluster space, with PEC subclusters 2 & 3 marking the root nodes. **(K)** Monocle3 pseudotime modules representing markers of the proliferative PEC and myofibroblast arms of previously established trajectory maps across PEC/myofibroblast/proliferative PEC clusters. **(L)** Heatmap of 19 Monocle3 pseudotime modules encompassing all PEC subclusters, myofibroblast and proliferative PEC clusters. **(M)** UMAPs of Monocle3 pseudotime modules chosen on the basis of discrete gene expression blocks across the PEC/Myofibroblast/Proliferative PEC cluster space. **(N)** WikiPathways analysis of Monocle3 pseudotime modules 2 & 4 - with trajectories oriented either towards the proliferative PEC cluster (Module 2) or towards the myofibroblast cluster (Module 4) from PEC subclusters.

### The transcription *factor Fosl2* has emerged as a central regulator of PEC activation and proliferation post-NTS

To better understand the potential mechanism(s) driving PEC activation, we first sought to identify early-inducible transcriptional regulators post-NTS in the PEC cluster. Peaks of accessible chromatin revealed a greater than 30-fold (log_10_/P-adj) enrichment of *Fosl2* transcription factor (TF) motifs (a member of the broader family of AP-1 immediate early genes) (30, 31) in PECs, when compared against all other clusters (**Figure 3A, Supplemental Table 5**). A ridge plot of *Fosl2* TF motif bias revealed a strong positive skewing, demonstrating that per-cell accessibility for this TF differs considerably from the average across all other clusters (**Figure 3B**). Mapping these TF bias results for the *Fosl2* motif onto a UMAP embedding of all NTS clusters further revealed the significant enrichment in both chromatin accessibility and gene expression of the *Fosl2* locus (**Figure 3C**). Additionally, PEC-specific snATAC-seq peak distribution in open chromatin was heavily biased towards intergenic and intronic locations, suggesting a regulatory/enhancer role for these PEC-enriched open chromatin sequences (**Figure 3D**). When we specifically interrogated the *Spp1* locus we unearthed six potential enhancer elements containing *Fosl2* motifs (**Figure 3E**). GO ontology analysis of PEC-enriched snATAC-seq peak locations revealed a marked preference for loci involved in cellular adhesion, membrane dynamics and chemotaxis (**Figure 3F**). To confirm induction of *Fosl2* occurs in PECs post-NTS, we conducted lineage tracing studies using pPEC.rtTA;TRE-Cre;eGFP mice, where PECs express eGFP. Immunofluorescence staining in NTS-treated *PEC-rttA^eGFP^* mice demonstrated co-expression of FRA2 (*Fosl2*), CD44, and eGFP in the glomeruli, validating the increase in FRA2/*Fosl2* in activated PECs (**Figure 3G**). Finally, we interrogated publicly available data from the Encyclopedia of DNA Elements (ENCODE) consortium (32) - specifically histone mark ChIP-seq data obtained from adult mouse kidney cortex. When we aligned our PEC-enriched ATAC-seq peaks with ENCODE histone mark coverage data, we obtained a clear enrichment of for PEC peaks at both promoter (H3K4me3 marked) and active enhancer (H3K27Ac + H3K4me1) elements (**Figure 3H**) (32). All together, these results suggest a potential regulatory role for *Fosl2* (AP-1 member) in the activation and subsequent proliferation of PECs post-NTS, through both gene-proximal (promoter) binding as well as gene-distal/intronic (enhancer) signatures.

**Figure 3:**
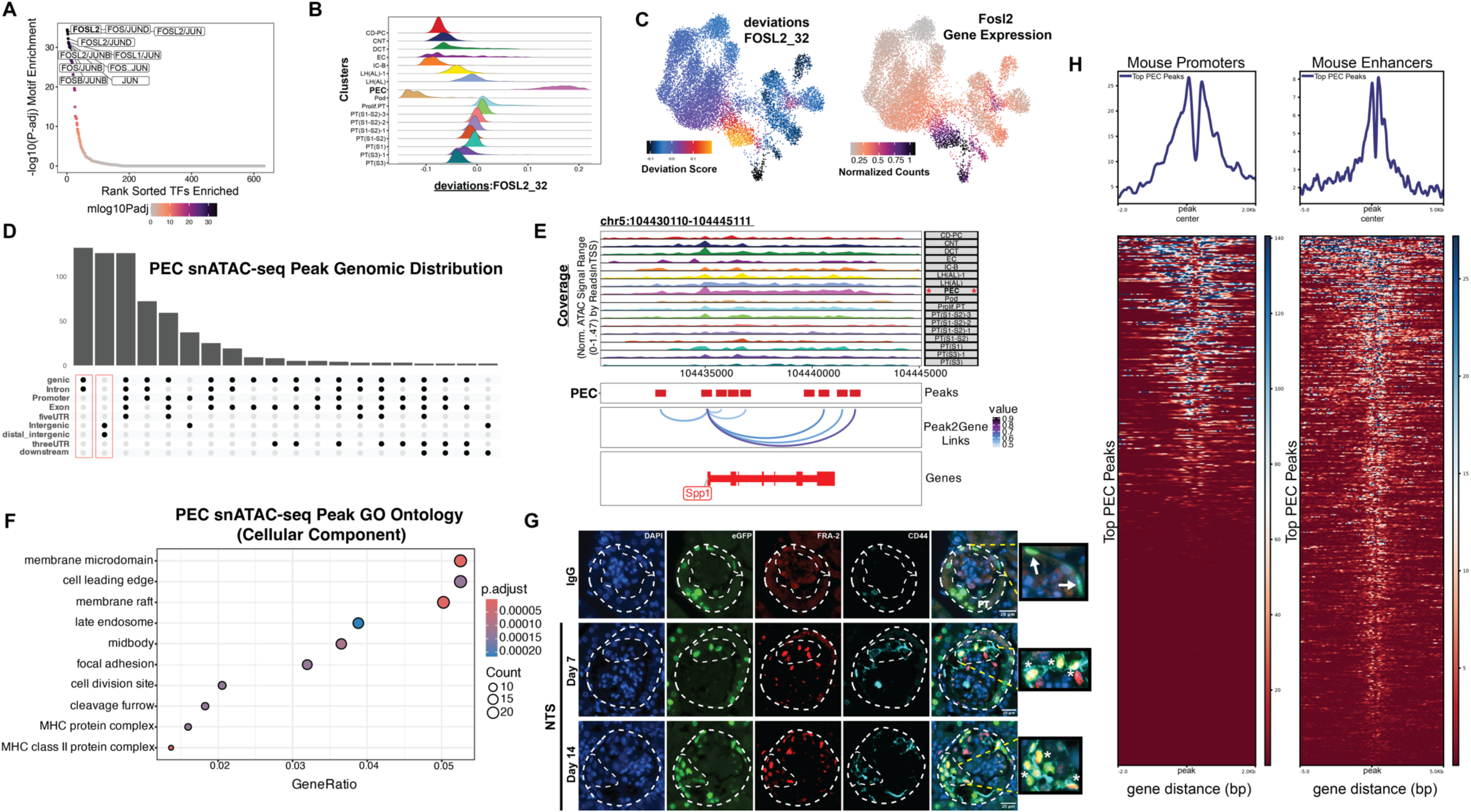
The transcription factor FOSL2 might regulate PEC activation and proliferation post-NTS treatment. **(A)** Rank sorted enrichment plot of transcription factor (TF) motifs enriched in the PEC cluster vs all other clusters in the NTS time-course. **(B)** ChromVAR deviations ridge plot of FOSL2_32 motif across all clusters. **(C)** ChromVAR FOSL2_32 motif deviation UMAP (left), *Fosl2* gene expression UMAP (right). **(D)** UpSet plot of the genomic distribution of PEC-specific ATAC-seq peaks. **(E)** ArchR peak-to-gene linkage genome cover plot of mouse *Spp1* locus. **(F)** GO Cellular Component ontology dot plot of enriched terms taken from list of genes proximal to PEC-specific ATAC-seq peaks. **(G)** Representative images of immunofluorescence staining in NTS-treated (day 7, 14) as compared to IgG-treated mice: eGFP (pPEC.rtTA - *hPODXL1*), FRA2 (*Fosl2*) & CD44. Arrows in IgG row point to eGFP+ quiescent PECs on Bowman’s capsule, and asterisks in day 7 & 14 NTS rows indicate triple colocalization (eGFP+/FRA2+/CD44+). Non-specific eGFP staining is noted in scattered PT cells. **(H)** DeepTools profile plots (top) and associated heatmaps (bottom) of top NTS multiome PEC-specific peaks mapped over the histone mark coverage signal of putative adult mouse kidney cortex gene promoter (H3K4me3) and enhancer (H3K27Ac + H3K4me1) elements obtained from the ENCODE consortium.

### FRA2/*Fosl2* protein-protein interactome displays enrichment of mRNA splicing factors and AP-1 heterodimer partners in cultured mouse PECs

Since we observed an enrichment of *Fosl2* motifs in PECs in the multiome analysis, we subsequently sought to characterize FRA2/*Fosl2* binding partners whose interactions might serve to regulate its function post-NTS. We initially employed Rapid Immunoprecipitation Mass Spectrometry of Endogenous Proteins (RIME) in cultured PECs to identify potential 1^st^ and 2^nd^ order binding partners (**Figure 4A**). Quantification revealed 31 high confidence candidate FRA2 interactors – with a minimum of 50% belonging to cellular pathways involved in mRNA processing and alternative splicing (**Figures 4B, C and Supplemental Tables 6 and 7**). Additionally, the RIME identified JUNB (established AP-1 heterodimerization partner of FRA2 (33, 34)), whose gene expression is also enriched in the PEC cluster (**Figures 4B, D**). Native immunoprecipitation for FRA2 demonstrated a robust pull-down of the JUNB protein, signaling a 1^st^ order interaction likely at the heterodimer interface in cultured PECs (**Figure 4E**). Interrogation of the top enriched FRA2-interacting proteins from the RIME (**Figure 4B)** in the PEC subclusters showed upregulation in PEC sub-clusters 1 & 2 (**Figure 4F**), with PEC sub-cluster 2 partially serving as the root for the pseudotime trajectory that eventually terminated in the downstream proliferative PEC/myofibroblast clusters (**Figure 2I**). Given that FRA2-interacting proteins are heavily involved in alternative splicing (**Figure 4C**), we employed the R package MARVEL to gain a better understanding of overall single-cell splicing dynamics in the NTS dataset (**Figure 4G**). “Coordinated” gene/splice junction (SJ) relationships were predominant in day 7 post-NTS during the putative transition from PECs to the myofibroblast cluster, while “isoform-switching” dynamics were observed primarily in day 14 post-NTS (**Figure 4G**). GO ontology performed on the genes undergoing *isoform-switching* revealed pathways involved in fibroblast proliferation, cell adhesion and RNA splicing (**Figure 4H**). Collectively, these data suggest a potential role for *Fosl2* in regulating alternative splicing and mRNA processing dynamics in PECs post-NTS, which might contribute to transition of PECs to myofibroblasts.

**Figure 4:**
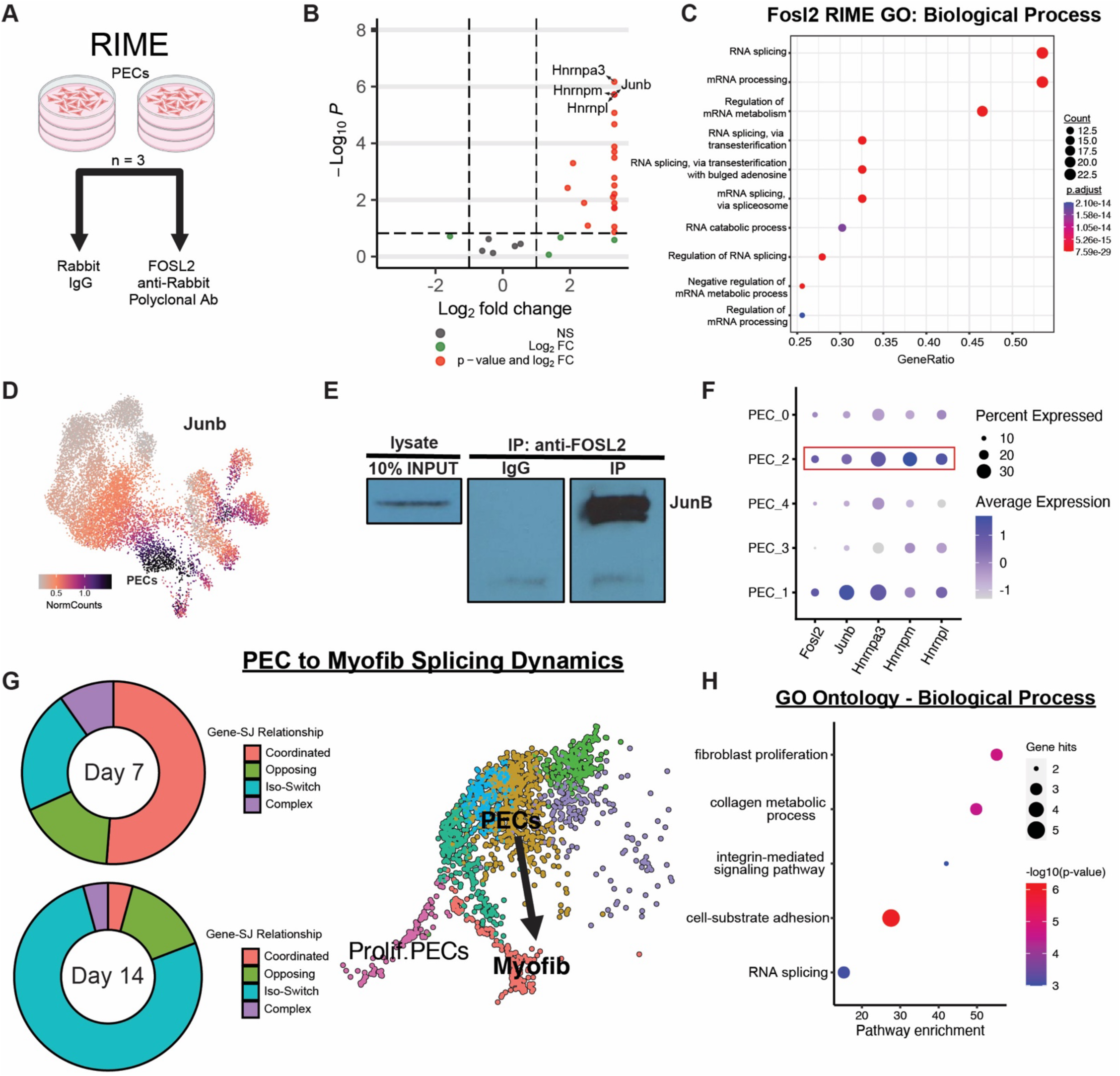
FRA2 (*Fosl2*) protein-protein interactome displays enrichment of mRNA splicing factors and AP-1 heterodimer partners in cultured PECs. **(A)** Schematic of RIME experiment, with n=3 per group 10cm plates of an immortalized mouse PEC cell line being employed to assay for FRA2 (*Fosl2*) interacting proteins/complexes (IgG vs FOSL2 anti-rabbit polyclonal antibody). **(B)** Volcano plot of IgG vs FRA2 (*Fosl2*) antibody RIME enrichment with log_10_ transformed p-values on the y-axis and log_2_ fold-change values on the x-axis. **(C)** GO Biological Process ontology dot plot of enriched terms derived from list of genes underlying proteins identified via FRA2 (*Fosl2*) RIME. (D) *Junb* UMAP feature plot embedding displaying PEC-specific enrichment of this top RIME hit and putative FRA2 (*Fosl2*) heterodimer partner. **(E)** Co-immunoprecipitation (Co-IP) of FRA2 (*Fosl2*) binding partners, blotting the 10% input lysate for FRA2 (*Fosl2*) (left), IgG control (middle) and JUNB target putative heterodimer binding partner (right). **(F)** Seurat DotPlot of the following RIME-enriched genes/proteins in PEC subclusters: *Fosl2*, *Junb*, *Hnrnpa3*, *Hnrnpm*, *Hnrnpl*. **(G)** MARVEL pie charts of Gene-SJ relationship proportions and UMAP between days 7 & 14 post-NTS: PECs, proliferating and myofibroblasts. Arrow indicates the splicing transition, specifically of the Iso-Switch category, across the PEC/Myofibroblast lineage boundary between days 7 & 14 of the NTS time course. **(H)** GO Ontology (Biological Process) of genes that undergo isoform switching between days 7 & 14 of the NTS time-course as determined by MARVEL analysis.

### Genetic lineage tracing of PECs post-NTS uncovers myofibroblast trans-differentiation events and CRISPR-Cas9 ablation of *Fosl2* in cultured PECs reduces PEC proliferation and expression of activated PEC/myofibroblasts markers

Given our NTS data of pseudotime bifurcation of PEC to proliferative PEC/myofibroblast cell states, we mapped the distribution of proliferation (*Mki67+*) and myofibroblast (*Acta2*) markers in blended UMAP space, clearly demonstrating separable gene expression signatures downstream of PEC activation (**Figure 5A**). This finding is supported by the presence of largely non-overlapping populations of KI67+ and alpha-smooth muscle actin (α-SMA)+ cells within NTS glomerular lesions, with each displaying a near complete overlap with the eGFP signal of lineage-traced PEC populations (eGFP+) (**Figure 5A**). Additionally, the eGFP+ activated PEC lineage displays a mixture of cells expressing α-SMA and FRA2 (*Fosl2*) - corroborating our bifurcating activated PEC cluster results from **Figures 2 & 5B**. To test the role of FRA2 (*Fosl2*) in PECs, we knocked-out *Fosl2* in cultured PECs (**Figure 5C**). We observed a significant decrease in proliferation in *Fosl2*-CRISPR KO clones as compared to Cas9-only control PECs (**Figure 5D**). Furthermore, α-SMA expression was significantly decreased in *Fosl2*-CRISPR KO clones as compared to Cas9-only control PECs - as was the activated PEC marker vascular cell adhesion molecule 1 (VCAM1) (**Figure 5E**). These data suggest that knockout of *Fosl2* in PECs reduces the expression of activated PEC and myofibroblast markers, while simultaneously reducing PEC proliferative capacity - pointing to its critical role in these cell states.

**Figure 5:**
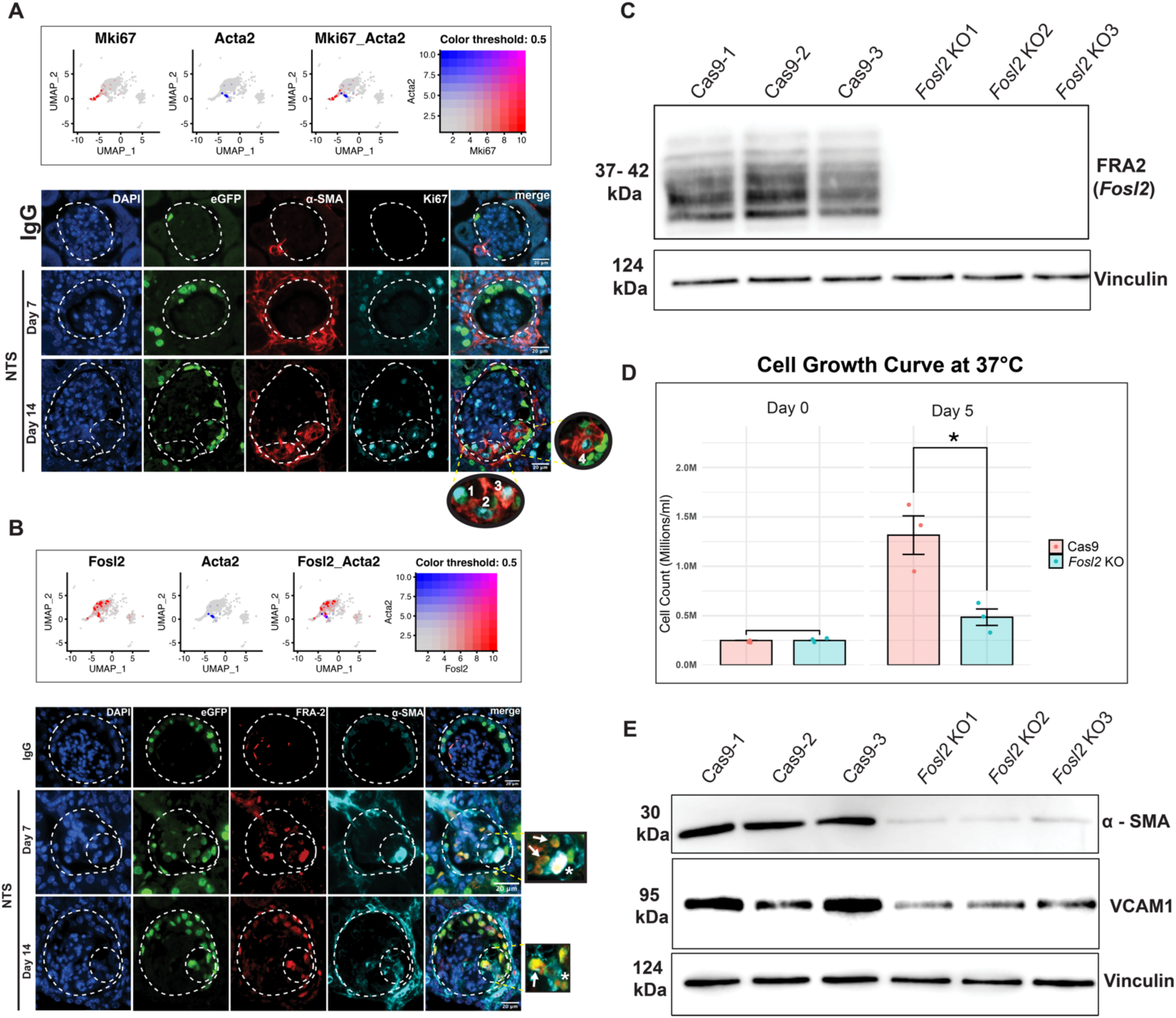
Genetic lineage tracing of subset of PECs to myofibroblast population post-NTS and consequences of CRISPR deletion of *Fosl2* in cultured PECs. **(A)** Top panel shows blended UMAP plots of PEC subclusters for *Mki67* and *Acta2*. Bottom panel shows immunofluorescence staining in IgG as compared to NTS-treated (day 7, 14) mice: eGFP (pPEC.rtTA - *hPODXL1*), α-SMA (*Acta2*), Ki-67 (*Mki-67*). Numerical indicators of potential PEC lineage crescentic cell-types: 1) Ki67+, 2) α-SMA+, 3) α-SMA+/Ki67+ and, 4) eGFP+ only **(B)** Top panel shows blended UMAP plots from PEC subclusters for *Fosl2* and *Acta2.* Bottom panel shows immunofluorescence staining for PECs/Myofibroblast in IgG as compared to NTS-treated (day 7, 14) mice: eGFP (pPEC.rtTA - *hPODXL1*), FRA2 (*Fosl2*), α-SMA (*Acta2*). Day 7 arrows indicate FRA2+ (*Fosl2*)/eGFP+ cells and the asterisk points to an example of PEC mid-trans differentiation into myofibroblast (eGFP+/FRA2+/α-SMA+). Day 14 asterisk points to pre transdifferentiating PEC marked with eGFP and FRA2 only. **(C)** Western blot for FRA2 (*Fosl2*) abundance in Cas9 only controls and *Fosl2* CRISPR-Cas9 deletion samples (n=3 each). **(D)** Bar plot of cell count differences between days 0 and 5 of cultured PEC cell growth in the context of *Fosl2* CRISPR-Cas9 deletion (n=3 each, Statistical significance between Cas9 and F1 genotypes at each time point was assessed using unpaired, two-tailed Welch’s t-tests. Comparisons were performed independently for Day 0 and Day 5 using three biological replicates per group. Data are presented as mean ± SEM, with individual data points shown). **(E)** Western blots from CRISPR-Cas9 *Fosl2* deletion in cultured PECs of canonical myofibroblast marker α-SMA and the activated PEC marker VCAM1, with housekeeping control vinculin (n=3 each).

### The PEC transcriptome in the *Klf4*^Δ*Pod*^ mice resembles the nephrotoxic serum-nephritis model

To validate the findings observed in the PECs post-NTS, we utilized a progressive murine model of FSGS with PEC activation and proliferation, podocyte-specific *Klf4* knockdown (*Klf4*^Δ^*^Pod^*) mice, where these mice develop severe albuminuria starting at 7-8 weeks of age with podocyte loss and subsequent PEC activation and proliferation, leading to FSGS (15). We performed renewed clustering on the snRNA-seq previously reported on *Klf4*^Δ^*^Pod^* and *Klf4^fl/fl^* kidney cortex (14, 15). Initial re-clustering yielded 18 clusters, which included the previously reported PEC cluster (**Supplemental Figures 10A-E,11A**). The enrichment of PEC activation marker *Cd44*, confirmed these were activated PECs (**Supplemental Figure 11B, Supplemental Table 8**). Sub-clustering the entire PEC population yielded 5 distinct sub-clusters, which resembled the sub-clustering from the NTS-nephritis model (**Supplemental Figure 11C**). The subcluster space displayed features strongly conserved across the NTS dataset, such as two sub-groups (PEC_2 & PEC_3) with a robust *Spp1/Cd44* co-expression signature (**Supplemental Figure 11D, Supplemental Table 9**). In addition, similar to PEC subclusters from the NTS model, PEC subclusters 2 & 3 revealed categories such as mRNA processing, focal adhesion and primary FSGS (**Supplemental Figures 11E,F**). Given the close apposition between mesangial cell 2 (MC2) and the PEC cluster, we subset all PEC subclusters with the addition of MC2 and observed that PEC_2 & PEC_3 phenocopied the PEC subclusters from the NTS-treated mice, wherein we could posit a signaling module involving *Spp1* and *Cd44* (**Supplemental Figure 11G**). An additional input could also be observed with the secretion of *Spp1* ligand by the MC2 cells, with PEC subclusters 2-4 serving as potential recipients via high *Cd44* expression (**Supplemental Figure 11H**). Subsequent pseudotime trajectory analysis showed root cells in PEC_2 & PEC_3 influencing downstream gene expression in a newly formed bridge area between MC2 and PEC_3 – which only appeared in the *Klf4*^ΔPod^ mice as compared to the NTS-nephritis model (**Supplemental Figure 11I**). However, gene expression modules 3 & 5 recapitulated the pseudo-time modules from the PEC subclusters post-NTS (**Supplemental Figure 11J**), which also yielded similarly enriched pathways involving primary FSGS and RHO GTPase signaling (**Supplemental Figure 11K**). Immunofluorescence staining for OPN (*Spp1*) and CD44 showed increased glomerular expression and colocalization of OPN (*Spp1*) and CD44 in SSeCKS+ (*Akap12*) PECs from *Klf4*^ΔPod^ mice as compared to *Klf4*^fl/fl^ mice (**Supplemental Figure 11L**). Conserved gene expression features in PECs between the NTS and *Klf4*^ΔPod^ models suggest shared signaling pathways during the development and progression of FSGS.

### *Fosl2* is enriched in PECs in both nephrotoxic serum-nephritis and *Klf4*^ΔPod^ murine models

We conducted *de novo* snATAC-seq in *Klf4*^ΔPod^ and *Klf4*^fl/fl^ mice to assess whether changes in chromatin accessibility in PECs were also similarly conserved with the NTS-nephritis model (**Supplemental Figure 12A-F**). Similar to post-NTS, multiple AP-1 subunit motifs, including *Fosl2,* were uniquely enriched in the PEC cluster (nearly 1K-fold) as compared to all other clusters (**Supplemental Figure 13A, B and Supplemental Table 10**). In addition to the PEC-specific *Fosl2* motif enrichment in the *Klf4*^ΔPod^ mice, its gene expression signature was also specific to the PEC cluster (**Supplemental Figure 13C**). To validate the likelihood of *Fosl2* sequence-specific chromatin binding in PECs, we utilized TF footprinting to demonstrate significant cut-site bias around the PEC *Fosl2* motif positions – as compared with podocytes which is another *Fosl2*-enriched disease cluster (**Supplemental Figure 13D**). We subsequently queried the array of PEC-enriched TFs for potential positive regulators, which are defined as those TFs whose open chromatin dynamics positively correlates with their own gene expression. *Fosl2* was identified as a significant positive regulator in PECs, further underscoring its critical role in PEC activation (**Supplemental Figure 13E**). Immunofluorescence staining for FRA2/*Fosl2* and CD44, validated the increase in FRA2/*Fosl2* in activated PECs (**Supplemental Figure 13F**) Collectively, similar to NTS nephritis model, these data suggest that there is an enrichment of FRA2/*Fosl2* in activated PECs in the *Klf4*^ΔPod^ mice, which might contribute to the progression of FSGS.

### FRA2 (*Fosl2*) genomic binding and expression in human kidney disease biopsies

We queried data from the ENCODE consortium (32), specifically histone mark ChIP-seq data from adult human kidney cortex. After mm10 (mouse) to hg38 (human) assembly genomic site lift-over and alignment of our PEC-enriched ATAC-seq peaks with ENCODE histone mark coverage data, we can demonstrate a robust enrichment of PEC peak abundance at both promoter (*H3K4me3* marked) and active enhancer (*H3K27Ac* + *H3K4me1*) elements (**Figure 6A**). In addition to the ENCODE histone ChIP-seq promoter/enhancer analysis, we wanted to assess the potential overlap between our NTS multiome PEC-enriched ATAC-seq peaks and previously published human *FOSL2* ChIP-seq datasets. To that end we chose a recent dataset (35, 36), focused on studies of human T helper cells (Th17), largely because their *FOSL2* interactome data strongly overlapped with our cultured PEC RIME analysis, specifically at the level of genes/proteins involved in mRNA processing and alternative splicing. PEC-enriched open chromatin peaks from our NTS multiome experiments showed a strong overlap with actual *FOSL2* binding data from the aforementioned Th17 studies, with the control CNT cluster peaks which are largely devoid of *Fosl2* binding motifs showing little to no enrichment. As a secondary control we produced randomly shuffled genomic locations of our mouse PEC peaks, which also did not show any overlaps with the Th17 *FOSL2* binding studies (**Figure 6B**). In addition to performing these comparative studies with human Th17 cells, we also obtained similar *FOSL2* ChIP-seq coverage data of two human epithelial cell lines (MCF7 & HepG2) from the ENCODE consortium (32) - which corroborated our Th17 findings (**Supplemental Figures 14A,B).** Upon plotting this log2 gene expression data against estimated glomerular filtration rate (eGFR) values obtained from previously reported expression arrays in micro-dissected glomeruli in *Nephroseq* (37) we observed an inverse correlation, with lower eGFR correlating with increased expression of these FOSL2-mediated genes (**Figure 6C**). To further validate these findings, we performed deconvolution analysis *(MuSIC2)*, which leveraged bulk RNA-seq data from 25 kidney biopsies with FSGS and 8 healthy control kidney tissues from donor nephrectomies (NEPTUNE cohort) to confirm the presence of activated PECs. Top cell-type markers for both PEC and downstream myofibroblast clusters from our NTS dataset were used to deconvolve data to reveal dominant PEC/fibroblast cell-type proportions in this NEPTUNE cohort (**Figure 6D**). Finally, immunofluorescence staining of human kidney biopsies from patients with Pauci-Immune Focal Crescentic GN (ANCA-associated) and Anti-Glomerular Basement Membrane disease (Anti-GBM) - demonstrated a clear increase in FRA2/*Fosl2* expression in Claudin-1 (CLDN1) positive PECs. (**Figure 6E**). % crescentic burden measures from these human kidney biopsies (**Supplemental Table 11**) showed a significant positive correlation with the presence of FRA2+/CLDN1+ activated PECs across sampled glomeruli suggesting a potentially vital role for FRA2 (*FOSL2*) in the acute stages of disease pathogenesis (**Figure 6F**). These data support a role for increased FRA2*/FOSL2* expression in activated PECs from subtypes of glomerulonephritis as is demonstrated in our schematic of PEC behavior underscoring a critical role for FRA2 (*FOSL2*) in triggering PEC activation and the subsequent transition to myofibroblasts (**Figure 6G**).

**Figure 6:**
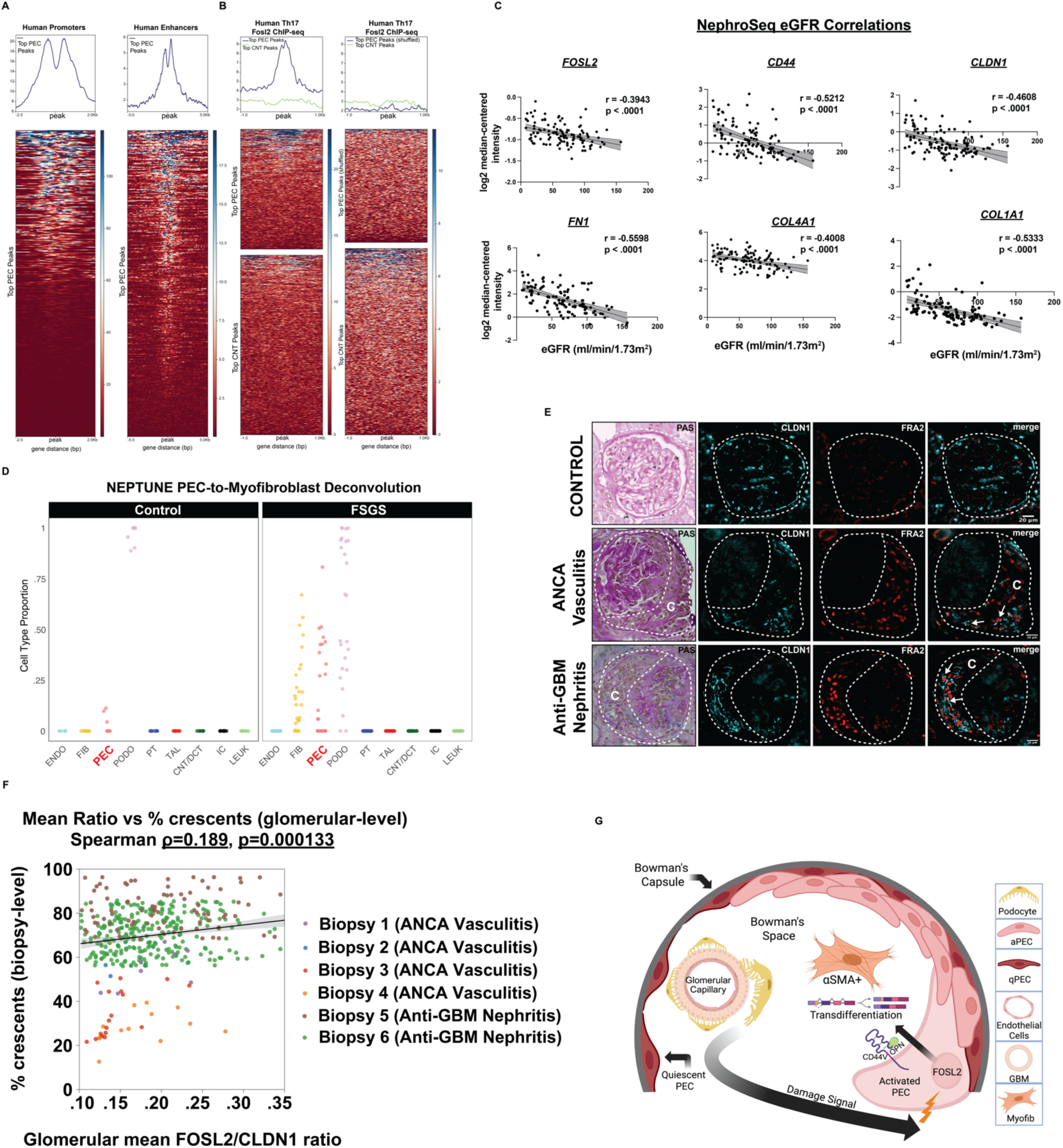
Mouse PEC open chromatin sites overlap with human kidney promoters/enhancers and Th17 FOSL2 ChIP-seq binding sites, with PEC-enriched gene expression negatively correlated with eGFR. **(A)** DeepTools profile plots (top) and associated heatmaps (bottom) of top NTS multiome mm10-to-hg28 lifted over PEC-enriched peaks mapped over the histone mark coverage signal of putative adult human kidney cortex gene promoter (H3K4me3) and enhancer (H3K27Ac + H3K4me1) ENCODE elements. **(B)** DeepTools profile plots (top) and associated heatmaps (bottom) of top NTS multiome mm10-to-hg28 lifted over PEC-enriched and CNT-enriched peaks mapped over the human Th17 *Fosl2* ChIP-seq (35, 36) coverage (left), with the right plot introducing randomly shuffled PEC-enriched peak data **(C)** Nephroseq Log2 expression plots of FOSL2 mediated genes (MuSIC2 deconvoluted genes) from the Ju CKD dataset (glomerular fraction) (37) against eGFR (mL/min/1.73m^2^). Scatter plot with linear regression employing eGFR values (x-axis) versus Log2 expression of FOSL2-mediated genes (y-axis). **(D)** Multi-subject Single-cell Deconvolution (MuSIC2) of FSGS bulk RNA-seq samples from 25 human kidney biopsies with FSGS and 8 control donor nephrectomies from the NEPTUNE cohort (74), using cell-type markers from our multiome snRNA-seq PEC and myofibroblast clusters. **(E)** Representative immunofluorescence/PAS staining of (*FOSL2* & *CLDN1*) from human kidney biopsies from donor nephrectomies as compared to kidney biopsies with ANCA vasculitis and Anti-Glomerular Basement Membrane (Anti-GBM) disease. The dotted line demarcates the human glomeruli with white arrows indicating sites of overlap between *FOSL2* and *CLDN1* - the latter being a PEC cell-type marker in both mice and humans. The dotted lines running through the middle of the glomeruli separate tuft from crescentic lesions (marked with large C). **(F)** Glomerular-level Spearman correlations across human kidney biopsies with ANCA vasculitis and Anti-GBM nephritis of FRA2+/CLDN1+ cells plotted against % glomerular crescent burden. **(G)** Proposed schematic of *Fosl2*/FRA2 in PEC alternative splicing and trans-differentiation to myofibroblasts.

## DISCUSSION

Mechanisms mediating PEC activation and subsequent transition to glomerulosclerosis remain poorly understood. Here, we utilize a combination of snRNA-seq and snATAC-seq in two independent murine models (transient and progressive) of glomerulosclerosis with PEC activation and proliferation to identify novel FRA2/Fosl2-mediated Spp1/Cd44 signaling in activated PECs. We also report that FRA2/Fosl2 potentially regulates alternative splicing in PECs, which might contribute to the activation of PECs into a “transition” state post-injury. Finally, we observed that the loss of *Fosl2 in-vitro* contributes to the blunted differentiation of activated PECs into myofibroblasts and lineage tracing studies *in-vivo* confirmed the transition of PECs to myofibroblasts post-NTS.

Our studies demonstrate a *Spp1*/Cd44 PEC-specific autocrine ligand/receptor loop, with *Spp1* expression preceding that of *Cd44* in the NTS time course (**Figure 1E**). A concordant rat-based NTS animal model similarly demonstrated an early rise in OPN/*Spp1* levels as well as co-localization of OPN and CD44 in crescent-associated PECs during disease progression (18), which we observed at later time points post-NTS. Additionally, digital spatial profiling of individual glomeruli in kidney biopsies with ANCA-associated glomerulonephritis demonstrated a strong correlation between *Spp1* and *Cd44* expression levels in PECs (38). Finally, our previous work implicated integrin signaling in PEC activation (15) which conforms with published data that outside-in integrin signaling through OPN (*Spp1*) and variable *Cd44V* isoforms plays a role in survival and apoptosis pathways - and may provide a mechanistic framework for further investigating OPN/CD44 signaling cascades in activated PECs (19).

Prior to the utilization of snRNA/ATAC sequencing approaches, PECs were treated as a monolith (39). Given their sparsity in non-pathological states, purifying pure cell populations for bulk RNA sequencing has been challenging (40). Recent studies of PEC molecular dynamics have leveraged public single-cell/nuclear RNA sequencing datasets - revealing, for the first time, some of the innate substructure present in pathogenic PEC subgroups (40). Although we observed similarities between these recent PEC subgroup breakdowns and our PEC subcluster data, especially in terms of similar canonical *Cd44+* proliferative activated PEC subclusters, our data did not point to any PEC subgroup containing the features of podocyte progenitors as outlined in the recent work by Liu et al. (40, 41). The addition of snATAC-seq in two independent murine models provides an inferential edge to previous work by Liu et al. (40) by enabling the characterization of PEC sub-populations at the gene regulatory level - as evidenced by the discovery of the role FRA2 (*Fosl2*) plays in activated PEC transcriptional and chromatin dynamics *in-vivo*. Sub-clustering of the PEC cluster in the NTS model revealed five subgroups, two of which exhibited strong co-expression of the previously described *Spp1*/*Cd44* ligand-receptor pair (**Figure 2D-G**). Pseudotemporal ordering starting within PEC subclusters revealed a strong PEC trajectory toward proliferative PECs/myofibroblasts, suggesting signaling pathways tied to primary FSGS, focal adhesion and glutathione metabolism (**Figure 2K-N**).

*Fosl2* expression was significantly increased in PECs, concordant with increased accessibility of its genomic consensus binding motif in both models. With prior evidence suggesting that *Fosl2* might be a critical regulator of PEC activation, we aimed to understand the FRA2/*Fosl2* protein-protein interactome. RIME, a method of immunoprecipitation followed by mass spectrometry - specifically designed to study chromatin and TF complexes suggests that FRA2/*Fosl2* interacts with splicing factors and RNA binding proteins. Although the breadth of AP-1 involvement in the splicing machinery has hitherto been largely unexplored, several studies have given credence to this potential connection. One such early study established a connection between T-cell MEK-ERK signaling and retention/displacement of variable CD44 exons and given that ERK signaling is a central upstream regulator of AP-1 activity we can posit the existence of potential co-transcriptional coupling between FRA2 (*Fosl2*) and the splicing machinery (42). This possibility is particularly attractive because we have observed a nearly complete elimination of ERK1/2 phosphorylation upon CRISPR-based deletion of *Fosl2* in cultured mouse PECs (data not shown), pointing to potential AP-1 feedback on the MEK-ERK pathway. Additionally, our RIME experiments revealed the potential interaction between FRA2 (*Fosl2*) and Sam68 (*Khdrbs1*) which is a core alternative splicing regulator (43–45), an interaction which we validated through native Co-IP experiments that demonstrated an RNA-independent interaction between FRA2 (*Fosl2*) and Sam68 (data not shown). This is quite interesting because of the established interaction between nuclear factor kappa-light-chain-enhancer of activated B cells (NF-Kb) family member p65 (*Rela*) and Sam68 splicing functions - given that *Rela* is also a highly enriched gene in activated PECs post-NTS induction and podocyte *Klf4* deletion (43, 46). These data support the potential connection between AP-1 and gene splicing, corroborated in a recent study of the *Fosl1* and *Fosl2* protein interactomes of T-helper 17 (Th17) cells, with raw *FOSL2* ChIP-seq data from these studies highlighting the overlap with mouse *Fosl2* motif-containing snATAC-seq PEC-specific open chromatin peaks (**Figure 6B**) (35). We confirmed through RIME and validated with native Co-IP that JUNB cooperates with FRA2/*Fosl2* to form the AP-1 heterodimerization subunit in activated PECs (**Figures 4D,E**). All in all, our data has revealed a previously unappreciated connection between AP-1 member FRA2 (*Fosl2*) and activated PEC dynamics in mouse models of crescentic glomerulonephritis through two potentially overlapping routes: 1.) Co-transcriptional coupling of AP-1 functions and its putative interaction with the core alternative splicing machinery in activated PECs, and 2.) The trans-differentiation of portions of the activated PEC cellular milieu into the α- SMA+ myofibroblasts which represents a novel route to the establishment of fibrotic/sclerotic lesions in affected glomeruli.

## METHODS

### Animal Experiments

The nephrotoxic serum (NTS) murine model in these studies utilized 8-week-old wild-type male mice on an *FVB/N* background as previously described (15). Briefly, 5 days prior to NTS administration, mice were pre-conditioned through the intraperitoneal (IP) injection of 0.5mg of Sheep IgG (Jackson ImmunoResearch) dissolved in Freund’s complete adjuvant (Millipore Sigma). Five days later at time point zero, all mice were administered 100ul of NTS or IgG control through an IP injection (12, 13, 15, 47). Urine was collected at regular intervals, specifically before the NTS injections, and subsequently at day 7 and 14 of the experimental time course. Mice were euthanized and transcardially perfused at Day 7 and Day 14 post-NTS injection – with kidney cortex samples harvested at all timepoints for histology, qRT-PCR and 10X Multiome analysis. *Klf4*^⊗^*^Pod^* genetic murine model of FSGS with PEC activation and proliferation on the *FVB/N* background was previously reported (15). Kidney cortex from 8-week-old *Klf4*^⊗^*^Pod^* and *Klf4^fl/fl^* male mice were used for snRNA/ATAC-seq experiments. Lineage tracing studies in the NTS model were performed using previously reported pPEC.rtTA (*hPODXL1*);TRE-Cre;eGFP mice (48).

### Study Approvals

The animal facility (DLAR) at Stony Brook Medicine is an AAALAC accredited institution which provides for strict control and management of any animal welfare issues. The DLAR has on its staff an animal health coordinator, animal technicians, and nurses. The staff monitored all animals daily and the Veterinarian was consulted if any animals appeared ill. A combination of ketamine and xylazine is employed via IP injection during mouse perfusions to minimize pain and reduce respiratory distress. Experimental protocols were all conducted in accordance with NIH Guide for the Care and Use of Laboratory Animals - as approved by the Stony Brook University Institutional Animal Care and Use Committee (IACUC). The rationale for the exclusive use to male mice in this set of studies stems from previously described sexual dimorphisms in a similar model that utilized a slightly different NTS formulation (Probetex, San Antonio, USA, PTX-001S lot#199-8)(49), and published accounts of sex differences in crescent development and mortality in lupus nephritis patients (50). Deidentified human kidney biopsies from University of Utah were scored for fibrosis by a renal pathologist (M.P.R). Control kidney biopsy specimens were acquired from healthy donor nephrectomies. The study was approved by the Stony Brook University Institutional Review Board.

### Cell Culture

Immortalized mouse PECs were used for all cell culture experiments in manuscript, as previously described (51). Briefly, cells were grown in <u>R</u>oswell <u>P</u>ark <u>M</u>emorial Institute (RPMI 1640) media supplemented with 10% fetal bovine serum (FBS), penicillin/streptomycin and Insulin-Transferrin-Sodium Selenite (ITS – Millipore Sigma). Immortalized Human Kidney Embryonic (HEK293T) cells were grown in Dulbecco’s Modified Eagle Medium (DMEM) with 10% fetal bovine serum (FBS) and penicillin/streptomycin.

### RIME (<u>R</u>apid Immunoprecipitation <u>M</u>ass <u>S</u>pectrometry of <u>E</u>ndogenous proteins)

Protocol was deployed with three biological replicate groups of both *Fosl2* shRNA target (N=3) and scramble control (N=3) cells. RIME was performed as previously described (20). FRA2 primary antibody and whole rabbit IgG **(Supplemental Table 12)** were used for the target and control plates, respectively. RIME samples were analyzed via LC-MS/MS with post-processing through Proteome Discoverer 2.4 **(Supplemental Tables 6,7)**. Immunoprecipitated samples were isolated on magnetic protein G beads, washed, and beads suspended in 100ul 5% SDS, 100mM triethyl ammonium bicarbonate (TEAB) for 5 minutes at RT. Protein-G beads were removed magnetically, and protein samples reduced by addition of dithiothreitol (DTT) to a final concentration of 10mM and incubated at 55°C for 30 min. Proteins were alkylated in 25mM iodoacetamide (IAc) at RT, for 30 min in the dark. The samples were digested with trypsin (20 ug) in 50 mM TEAB in a humified incubator overnight at 37 degC. Peptides were eluted by sequential addition of 80ul 50 mM TEAB, 0.2% formic acid, and 50% acetonitrile, 0.2% formic acid, each followed by centrifugation at 4000 x g for 1 minute. The samples were then dried by SpeedVac and resuspended in 0.1% formic acid (FA) and peptides analyzed by C18 reverse phase LC-MS/MS. HPLC C18 columns were prepared using a P-2000 CO2 laser puller (Sutter Instruments) and silica tubing (100µm ID x 20 cm) and were self-packed with 3u Reprosil resin. Peptide identification and quantitation was performed using an orbital trap (Q-Exactive HF; Thermo) instrument followed by protein database searching. Replicate samples were analyzed, using two different HPLC gradient profiles (0-30% ACN over 90’ and 0-40% ACN over 90’). Data files were acquired with Xcalibur. Peptide alignments and quantitation were performed using Proteome Discoverer v2.4 software (Thermo). Protein false discovery rates experiments are binned at 0.01 and 0.05 FDR. Peptide and PSM FDR cutoffs are typically set to 0.0. Label free quantitation (LFQ) was used for quantitation using pairwise peptide ratios for normalized abundance calculations. The mouse UniProt dataset (16982 entries) were used for data alignment. Matched peptide-based label free quantitation and p-values were calculated by Benjamini-Hochberg correction for FDR. Additional data sieving and creation of graphical representations were completed in RStudio using the EnhancedVolcano package.

### Histological Staining & Scoring

<u>P</u>eriodic <u>A</u>cid <u>S</u>chiff (PAS – Sigma Aldrich) staining was performed on Formalin-Fixed Paraffin-Embedded (FFPE) kidney tissue in the IgG control and day 7 & 14 post-NTS treated animals. Double-blinded semi-quantitative scoring of the following histopathological features were conducted in PAS-stained sections from NTS-treated mice and controls: % crescents, % FSGS lesions, % tubular injury and % interstitial inflammation.

### Single-nucleus isolation methods

The preparation of purified nuclei for all downstream applications such as 10X snRNA/ATAC-seq and 10X multiome was performed as previously outlined (52).

### Multiome, snRNA and snATAC-seq raw data processing

For samples from the NTS and *Klf4* datasets, demultiplexed fastq files from the Novaseq 6000 platform were aligned with the reference mouse genome (GRCm39) via 10X Cell Ranger Arc 2.0 (NTS), 10X Cell Ranger 7.0 (Klf4) and 10X Cell Ranger ATAC 1.2 (*Klf4*), using four compute nodes (running ten tasks per node) on a Cent-OS high performance cluster with a gene annotation general feature format (GTF) file in conjunction with the full genome sequence assembly fasta file. For the NTS dataset multiome STARsolo alignment, the Cell Ranger Arc whitelist file (737K-arc-v1.txt) was employed, and Cell Ranger 4-type adapters were clipped. For the *Klf4* independent snRNA and snATAC STARsolo alignment, standard Cell Ranger whitelists were used (53–55).

### Single-Cell Analysis of Regulatory Chromatin in R (ArchR)

Independent arrow files were made, using both the Cell Ranger-produced fragment file for both the NTS and *Klf4* datasets, with each group being merged into an ArchR project. Following doublet removal, snRNA-seq matrices were imported into the ArchR project. Iterative latent semantic indexing (LSI) was employed for the dimensionality reduction of both the snATAC and snRNA-seq components of the multiome and non-multiome datasets, with the latter involving constrained and unconstrained snRNA-seq integration steps. Uniform manifold approximation (UMAP), t-distributed stochastic neighbor embeddings (tSNE) and Harmony (batch effects corrected) embeddings were generated and used for most downstream purposes, with the exception of the Marvel splice junction analyses which utilized tSNE plots generated through SingCellaR (56). Model-based analysis of ChIP-seq (MACS2) was the snATAC-seq peak caller deployed through the ArchR R-Studio package. The remaining components of the ArchR analysis pipeline used as directed by the vignette: (https://www.archrproject.com/bookdown/index.html) - ChromVar motif deviation/enrichment, motif footprinting, peak2gene linkage and pseudotime trajectory analysis (57).

### Seurat (R toolkit for single cell genomics)

Seurat objects were created for both the NTS and *Klf4* datasets by reading in the 10X_h5 files into R Studio. For both datasets RNA features were subset to the range of 200-5000, with the mtDNA cutoff being set to a maximal value of 10%. SCTransform was employed for all individual Seurat objects prior to any merging of the individual datasets. Integration features were subsampled (2K) and SCT-prepped integration anchors were identified across all individually merged objects. Principal components analysis (PCA) and UMAP approaches were applied to all anchor integrated objects, with the assistance of elbow plotting to help identify the appropriate dimensionality of each merged dataset (e.g., # of clusters). All previously mentioned approaches utilized 1:30 dimensions, with the DimHeatmap function adding an additional layer of QC for assigning appropriate cluster resolution. Markers across all clusters were identified with SCT and RNA assay types. These top differential genes were then utilized for manual cluster cell-type annotation. Following the assignment of new cluster ids (specific kidney cell-types) to the Seurat object metadata, the following plotting functions were used to characterize the results: DotPlot, VlnPlot, RidgePlot, FeaturePlot. The FindSubCluster function was used for more refined clustering of the PEC groups in the NTS and *Klf4* datasets, subsets of which were subsequently saved as new Seurat objects. All data analysis was saved at both the level of R script and Seurat object specific RDS files, which were subsequently reloaded for continued analysis (58, 59).

### CellChat (Ligand-Receptor Interaction tool)

As input for ligand-receptor interaction analysis, for both the NTS and *Klf4* datasets, multiomic and separate/integrated ArchR objects (respectively) were used as inputs to create CellChat objects, with cell-type labels set as specific identifiers. The mouse CellChat database (http://www.cellchat.org/cellchatdb/) was set as the default list of reference ligand/receptor pairs. CellChatDB is split up into 3 categories: Secreted Signaling, ECM-Receptor Interactions and Cell-Cell Communication. All three categories were probed in the NTS and *Klf4* analytical pipelines. After computing the communication probabilities, all data points with fewer than 10 cells were removed from the downstream analysis. For the SPP1 pathway, which is highlighted in the manuscript, we used the following plotting types: chord diagram, scatter plots of incoming/outgoing interaction strengths (26).

### Salmon Alevin

Prior to all Salmon Alevin transcript quantification, we used the following bamtofastq tools (https://www.10xgenomics.com/support/software/cell-ranger/latest/miscellaneous/cr-bamtofastq) to convert Cell Ranger 7.0 generated BAM files back to fastq files. We next employed the newly generated read 1 and 2 fastq files along with a previously constructed Salmon mouse transcriptome index (generated using fasta file of GENCODE vM34), plus a .tsv file containing transcript ID to name mapping – to quantify isoform abundance in our 3’ tagged-end single-cell sequencing data. For each dataset the --forceCells flag was used along with the number of cells from Cell Ranger 7.0 alignment. In addition, the 10X chemistry used was denoted with the -- chromiumV3 flag (60).

### STARsolo (mapping, demultiplexing and quantification for single cell RNA-seq)

Before MARVEL analysis of splicing dynamics, we utilized STARsolo to obtain the unique molecular identifier (UMI) counts per cell barcode for the following categories under the --soloFeatures flag: Gene (counts that map to unique genes), GeneFull (counts mapping to introns and exons of all genes), SJ (splice junction counts), Velocyto (spliced, unspliced and ambiguous counts per gene/cell). The inputs for STARsolo mapping were all raw fastq files obtained from 150bp Illumina NovaSeq reads, which followed after either 10X Multiome (NTS) or *Klf4* (separate snRNA and snATAC runs) droplet based single-nuclear preparations. STARsolo runs consisted of individual jobs on two of the Stony Brook University 28-core high performance cluster nodes (53, 54).

### SingCellaR (R functions for scRNA-seq data)

Additionally, to set up MARVEL objects we employed this package for the generation of gene-level descriptors (count matrices, cell/gene metadata) for both the NTS and *Klf4* datasets. SingCellaR functions support multiple integrative approaches such as Seurat overlays. For the MARVEL input we only used the SingCellaR generated count matrix as well as phenoData (cell metadata) and featureData (gene metadata) (56).

### MARVEL (Single Cell Splicing Dynamics)

The MARVEL object was constructed out of three primary components: 1.) count matrix and cell/gene metadata at the filtered gene level (SingCellaR), the raw gene and the splice junction (SJ) level, 2.) a file containing UMAP coordinates based on the filtered gene metadata, 3.) and finally the GENCODE vM34 annotation .gtf file. Once created from all of these components, the MARVEL object is annotated, meaning that genes/SJs are assigned to type (e.g., protein-coding, lncRNA, miRNA etc.). Upon deciding which gene category to study (protein-coding), we looked specifically at the transition between PECs and myofibroblasts on days 7 and 14. We utilized circular bar plots to emphasize the shift in Gene-SJ relationship dynamics which has four possible categories (Coordinated, Opposing, Iso-Switch, Complex) based on the MARVEL integrative data analysis model. The varying gene sets underlying this newly revealed set of Gene-SJ dynamics were used as inputs for GO Ontology analysis (61).

### scVelo (RNA Velocity Analysis)

As a first step, ArchR objects were converted to SciPy AnnData objects though the SeuratDisk package in R (62). These initial objects were derived from Salmon Alevin output data which included information about the overall proportions of spliced vs unspliced transcripts in the dataset. scVelo was then utilized via Jupyter notebook to perform all RNA Velocity analyses. Correlation/trendlines in velocity plots of spliced/unspliced transcripts were determined to be present by scVelo (command: scv.pl.velocity) via the overall directionality trend, with a dotted diagonal line denoting the spliced/unspliced breakpoint (27, 63, 64).

### Monocle3

Monocle3 pseudotime trajectory analysis used SeuratWrappers to convert Seurat to Monocle3 objects. Pseudotime trajectory-oriented cell clustering, in addition to the native Seurat clusters present in the converted object, was performed via Monocle3 whereupon both cluster modalities can be plotted through UMAP representations. Following these initial steps, pseudotime trajectories were plotted based on the identification of root cells and directionality inference was color-coded atop the same UMAP – only biased by the location of the root cell input. Additionally, Monocle3 was used to identify trajectory and branchpoint specific gene modules whose composition underlies the pseudotemporal order of cells along branches of the trajectory tree. Each gene module was then used to enumerate biological pathway (WikiPathways) differences between branches of the pseudotemporal trajectory tree (65–67).

### GO Ontology and Pathway Analysis

For all the pathway analysis in manuscript, the following R packages were used: clusterProfiler (Version 4.14.3 - 2024), ReactomePA (Version 1.50.0 - 2024), org.Mm.eg.db. The following pathway database tools were used: REACTOME, WikiPathways and KEGG. Gene ontology analysis was performed through GO categories: Molecular Function, Biological Process and Cellular Component. Basic enrichment and Gene Set Enrichment Analysis (GSEA) was performed throughout the manuscript, with the latter analytical pipeline employing the Molecular Signatures Database (MSigDB). All of the above-mentioned biological pathway and gene ontology analyses were performed in RStudio (68–72).

### MuSIC2 (Cell Type Deconvolution for Multi-Condition Bulk RNA-seq Data)

For the healthy adult human kidney scRNA-seq reference we used GSE185948 (73), containing scRNA-seq profiles from 5 healthy controls. The bulk RNA-seq deconvolution NEPTUNE cohort (GSE197307) (74) contained 25 FSGS samples and those from 8 healthy living kidney donors.

### NephroSeq

RNA sequencing deposited in Nephroseq database from human micro-dissected glomeruli of kidney biopsies with diabetic kidney disease (DKD), FSGS, minimal change disease (MCD), Lupus Nephritis, IgA Nephropathy, renal vasculitis and healthy control specimens was utilized to interrogate potential correlations between estimated glomerular filtration rate (eGFR) and the glomerular expression of *FOSL2*.

### DeepTools

Re-analysis of publicly available histone and TF ChIP-seq datasets was performed by initially creating matrices of ChIP-seq BigWig coverage files intersected with BED genome interval files of peak locations drawn from our NTS multiome (ATAC-seq) experiments. In the case of the mouse histone mark matrices (<u>promoters</u>: H3K4me3 coverage, <u>enhancers</u>: intersection of H3k27Ac/H3K4me1 coverage) the peak locations identified in our NTS multiome experiments were used directly, while in the case of the human histone mark and TF DeepTools matrices - we used the UCSC liftOver utility to transpose the genomic peak locations from the mouse mm10 assembly into the human hg38 assembly, prior to generation of the matrix files. Matrices were generated with the computeMatrix command using a single reference point (ATAC-seq peak center) with binning set to one and genomic locations spanning from +/- 1kb around the peak center to +/- 5kb, depending on the lateral genomic spread of the ChIP-seq signal coverage. Finally, the DeepTools plotHeatmap function was employed to generate both the profile plots and underlying heatmaps for all of the histone mark and TF targets.

### Immunofluorescence

All specimen slides were incubated in a 55°C oven for one hour to initiate deparaffinization, as previously described (14, 15). Briefly, formalin-fixed sections following initial oven incubation underwent one further round of deparaffinization with Xylene, after which point they were placed in a pressure cooker to facilitate antigen retrieval in a 1X Sodium Citrate Buffer solution – where the internal pressure cooker temperature was held at 110°C for 10 minutes before a gradual cooling period and subsequent depressurization of the chamber. Permeabilization was then performed with 0.2% Triton X-100 (Sigma Aldrich) for 10 minutes. A solution of 2% non-fat milk used was dissolved in 1X <u>T</u>ris-<u>B</u>uffered <u>S</u>aline with <u>T</u>ween (TBS-T) in which the slides were incubated for 1 hour at 37°C to effectuate blocking of non-specific antibody binding. Primary antibodies **(Supplemental Table 12)** were then applied to the sections for overnight (O/N) incubation at 4°C. On the second day of the protocol primary antibodies were washed off with 3 successive washes in TBS-T and secondary antibodies **(Supplemental Table 12)** were applied and the sections were incubated at 37°C for 30 minutes. Following three additional TBS-T washes, DAPI nuclear counterstain was applied to the sections for 10 minutes at room temperature and the slides were cover-slipped and left to dry O/N in the dark at room temperature (R/T).

### FIJI-Based Quantification of FRA2⁺/CLDN1⁺ Perinuclear Signal

Quantification of FOSL2-positive nuclei and associated perinuclear CLDN1 signal was performed using a custom JavaScript macro executed in Fiji (ImageJ v1.54). Multichannel ND2/TIFF images were imported and automatically split into individual channels. The nuclear (FOSL2) channel underwent Gaussian smoothing (σ = 0.8), Otsu thresholding, hole-filling, morphological opening/closing, and watershed segmentation to isolate individual nuclei. Candidate nuclei were filtered based on minimum mean nuclear intensity. For each accepted nucleus, a peri-nuclear “ring” mask was generated by isotropically enlarging the nuclear ROI by 4 pixels and subtracting the original ROI boundary. The CLDN1 channel was corrected using rolling-ball background subtraction (radius = 30 px), and mean ring intensity was measured for every nucleus. For each nucleus, red (nuclear), cyan (perinuclear), and ratio (cyan/red) intensities were recorded. Images containing ROIs were saved as PNG overlays, and per-image CSV files were automatically generated. A batch-mode implementation allowed entire biopsy folders to be processed, producing both per-image CSVs and a master merged dataset per patient. Only nuclei meeting pre-specified red- and cyan-intensity thresholds were retained for downstream clinical correlation.

### Image/Clinical Data Correlation Analysis

Quantitative per-glomerulus staining measurements generated from Fiji analysis (mean nuclear FOSL2 intensity, mean perinuclear marker intensity, and their derived ratio) to correlate with percentage of crescents. To perform correlation, each glomerulus was preserved as an independent measurement and annotated by patient identifier, ensuring that multiple glomeruli from the same biopsy contributed proportionally to the analysis. All numeric variables were inspected for non-normality using descriptive distributions, after which non-parametric Spearman rank correlations were performed. For each clinical measure, glomerular-level staining metrics (including the FOSL2-to-marker signal ratio) were evaluated against the corresponding biopsy-level readout. Correlation coefficients (ρ) and exact two-tailed significance values were reported. For visualization, glomerulus-level scatter plots were generated using seaborn regression (Python), displaying individual glomeruli as datapoints alongside fitted trend lines, and annotated with computed Spearman statistics. Only glomeruli with complete paired staining-clinical data were included in each correlation analysis.

### Serum Creatinine Cleaning and eGFR Calculation

Serum creatinine values were parsed and converted to numeric format and eGFR was calculated using the race-free CKD-EPI creatinine equation (75), which incorporates sex-specific κ and α parameters, nonlinear creatinine scaling, and age adjustment.

### CRISPR-Cas9 Gene Editing

For generating *Fosl2* KO CRISPR clones we employed the lenti-CRISPRV2 system (76) in an immortalized mouse PEC cell line (51). Two independent sgRNAs both targeting exon 3 of the *Fosl2* gene were selected (Synthego sgRNA design tool) based on their specificity and reduced potential for off-target effects. After cloning the *Fosl2*-targeting sgRNAs into the lenti-CRISPRV2 backbone (Addgene #52961) we generated high-tighter competent lentiviral stocks in HEK293T cells and proceeded to infect our cultured PECs. Following puromycin selection, we employed FACS to sort single *Fosl2* CRISPR clones into a 96-well plate. Eight final clones were chosen based on the following criteria: 1) No protein expression of FRA2 (*Fosl2*) in PEC lysates, 2) No transcript expression via qRT-PCR readout and finally, 3) Pyrosequencing confirmed deletions present.

### Co-IP (Co-Immunoprecipitation)

A protocol from Cell Signaling technologies was used along with minor modifications. Briefly, 1X IP Lysis Buffer (Pierce) was added to cells in 10cm plates and they were incubated for 5 minutes on ice before being scraped off, sonicated, and centrifuged at 14,000xg for 10 minutes at 4°C. The supernatant was then directly utilized in Co-IP reactions. The lysate was incubated with 20μl of pre-washed Protein A paramagentic beads (Invitrogen) with rotation at room temperature. Following these pre-clearing steps, primary antibodies for FRA2 and JUNB (targets) and IgG (controls) were added and incubated overnight with the pre-cleared lysate at 4°C with rotation. On the second day of the protocol, the lysate/antibody complex was incubated for 20 minutes with a new pre-washed magnetic bead pellet. Following 5 washes with 3X SDS buffer, the sample was heated to 95-100°C for 5 minutes and the beads were separated away with a magnetic rack. Subsequent samples were used to probe for FRA2 (direct target) and JUNB (interacting target) using 10% input.

### Western Blots

Protein lysate supernatants dissolved in radioimmunoprecipitation assay buffer (RIPA) buffer were all obtained from an immortalized PEC cell line (51), in the presence of protease/phosphatase inhibitors. All antibodies used in western blot experiments are listed in **Supplemental Table 12.**

### RT-PCR & qPCR

RNA was obtained from 6-well plates growing immortalized PECs through TRIzol extraction. Cells were washed twice with 1X PBS after which 250ul TRIzol reagent was added per well. TRIzol cell extracts were subjected to a standard Chloroform:Isoamyl RNA extraction protocol and the RNA concentration was quantified via a NanoDrop spectrophotometer. Matching quantities of RNA (1ug) were reverse transcribed with the SuperScript VILO cDNA Synthesis Kit. cDNA at a 1:20 dilution was used for all qPCR (SYBR Green) and RT-PCR experiments. All primers used in qPCR/RT-PCR experiments are listed in **Supplemental Table 13.**

### Statistical Analysis

Sample sizes were based on effect sizes from prior *Klf4*^ΔPod^ studies (15). Data distribution was assessed for normality, with parametric tests (*t*-test, two-way ANOVA with Tukey post hoc) applied to normally distributed data and nonparametric tests (Mann–Whitney) used otherwise. Associations were evaluated by Pearson correlation and simple linear regression. Tests used are noted in figure legends. Data are mean ± SEM from ≥3 independent experiments, with *P* < 0.05 considered significant (GraphPad Prism v10.0). Relationships between double-positive PEC burden and each histopathologic or clinical variable (eGFR, % crescents, % fibrosis) were evaluated using: Pearson correlation coefficient (r), Spearman rank correlation coefficient (ρ), Two-tailed *p-values*. Pairwise deletion was used to handle missing values. Simple linear regression was used to evaluate directional associations between double-positive PECs ↔ % Crescents. Model were fit using *scikit-learn* (Linear Regression).

### Data and Code Availability

All raw transcriptomic and chromatin accessibility data (10X Multiome) have been deposited in the Sequence Read Archive (Accession #: PRJNA1338507), with proteomics datasets being uploaded to the MassIVE Database (accession #: MSV000099252). All code/scripts used for the purposes of data analysis in this manuscript can be found at: https://github.com/MallipattuLab/Fosl2-manuscript

## AUTHOR CONTRIBUTIONS

RB, YG, MPR, and JDH performed the experiments and analyzed the subsequent data. RB and JDH performed the computational analysis. FS, VD and MPR provided services related to analyzing both mouse and human renal histopathological samples. JDH performed the mass spectrometry and analysis of the RIME samples. DJS provided the NTS used in the animal studies. DJS and JCH provided overall guidance on construction and scope of the experiments. RB and SKM designed the overall studies, wrote and edited the manuscript. Guarantors of this work and, as such, had full access to all the data in the study and take responsibility for the integrity of the data and the accuracy of the data analysis: RB and SKM.

## ACKNOWLEDGEMENTS

None

## DATA SHARING STATEMENT

Sequencing data were deposited in Sequence Read Archive (Accession #: PRJNA1338507). Proteomics data were uploaded to the MassIVE Database (accession #: MSV000099252).

