## Supplemental Figures 1 - 14 for "*Fosl2* regulates the transition from parietal epithelial cells to myofibroblasts in the kidney"

### TABLE OF CONTENTS

1. **Supplemental Figures + Legends:**
   **Supplemental Figure 1:** 10X Multiome data quality control plots from the NTS model.
   **Supplemental Figure 2:** CellChat CD44 ligand/receptor maps from the NTS model.
   **Supplemental Figure 3:** 10X Multiome UMAPs for key PEC-enriched genes in the NTS model.
   **Supplemental Figure 4:** Gene ontology and KEGG pathway GSEA for PEC cluster using the multiome dataset in the NTS model.
   **Supplemental Figure 5:** 10X snRNA-seq data quality control plots from the NTS model.
   **Supplemental Figure 6:** KEGG pathway analysis of DEGs from the PEC sub-clusters in the NTS model.
   **Supplemental Figure 7:** REACTOME pathway analysis of DEGs from the NTS snRNA-seq PEC sub-clusters in the NTS model.
   **Supplemental Figure 8:** WikiPathway analysis of DEGs from the NTS snRNA-seq PEC sub-clusters in the NTS model.
   **Supplemental Figure 9:** Monocle3 top 50 PEC pseudotime modules in the NTS snRNA-seq dataset.
   **Supplemental Figure 10:** 10X snRNA-seq data quality control plots made in Seurat for $Klf4^{fl/fl}$ versus $Klf4^{\Delta Pod}$ *mice*.
   **Supplemental Figure 11:** 10X snRNA-seq data analysis (Seurat, CellChat, Monocle3) for $Klf4^{fl/fl}$ versus $Klf4^{\Delta Pod}$ *mice*.
   **Supplemental Figure 12:** 10X snATAC-seq data quality control plots made in ArchR for $Klf4^{fl/fl}$ versus $Klf4^{\Delta Pod}$ *mice*.
   **Supplemental Figure 13:** ArchR standard data analysis pipeline with immunofluorescence staining in $Klf4^{fl/fl}$ and $Klf4^{\otimes Pod}$ mice.
   **Supplemental Figure 14:** DeepTools profile plots and associated heatmaps of overlap between mouse NTS open chromatin peak locations and FOSL2 ChIP-seq binding sites in human epithelial cell lines.

2. **Supplemental Tables: Supplemental_Tables_all.xlsx**
   **Supplemental Table 1:** Histopathological scoring of NTS nephritis model.
   **Supplemental Table 2:** NTS multiome upregulated/downregulated DEGs.
   **Supplemental Table 3:** NTS snRNA-seq upregulated/downregulated DEGs.
   **Supplemental Table 4:** NTS snRNA-seq upregulated/downregulated DEGs (PEC sub-clusters vs. all other PEC sub-clusters).
   **Supplemental Table 5:** NTS snATAC-seq upregulated/downregulated accessible peaks.
   **Supplemental Table 6:** Cultured PEC cell *Fosl2* RIME LC-MS/MS Proteome Discoverer 2.4 output spreadsheet detailing unique peptide counts and PSMs.

**Supplemental Table 7:** Cultured PEC cell *Fosl2* RIME LC-MS/MS Proteome Discoverer 2.4 output spreadsheet detailing distribution of post-translational modifications.
**Supplemental Table 8:** *Klf4* snRNA-seq upregulated/downregulated DEGs.
**Supplemental Table 9:** *Klf4* snRNA-seq upregulated/downregulated DEGs (PEC sub-clusters vs. all other PEC sub-clusters).
**Supplemental Table 10:** *Klf4* snATAC-seq upregulated/downregulated accessible peaks.
**Supplemental Table 11:** Human biopsy clinical characteristics and histopathological scoring.
**Supplemental Table 12:** List of antibodies used for: Immunofluorescence, western blot, RIME and co-immunoprecipitation.
**Supplemental Table 13:** Primers for real-time PCR.

**SUPPLEMENTAL FIGURES**

**Supplemental Figure 1**

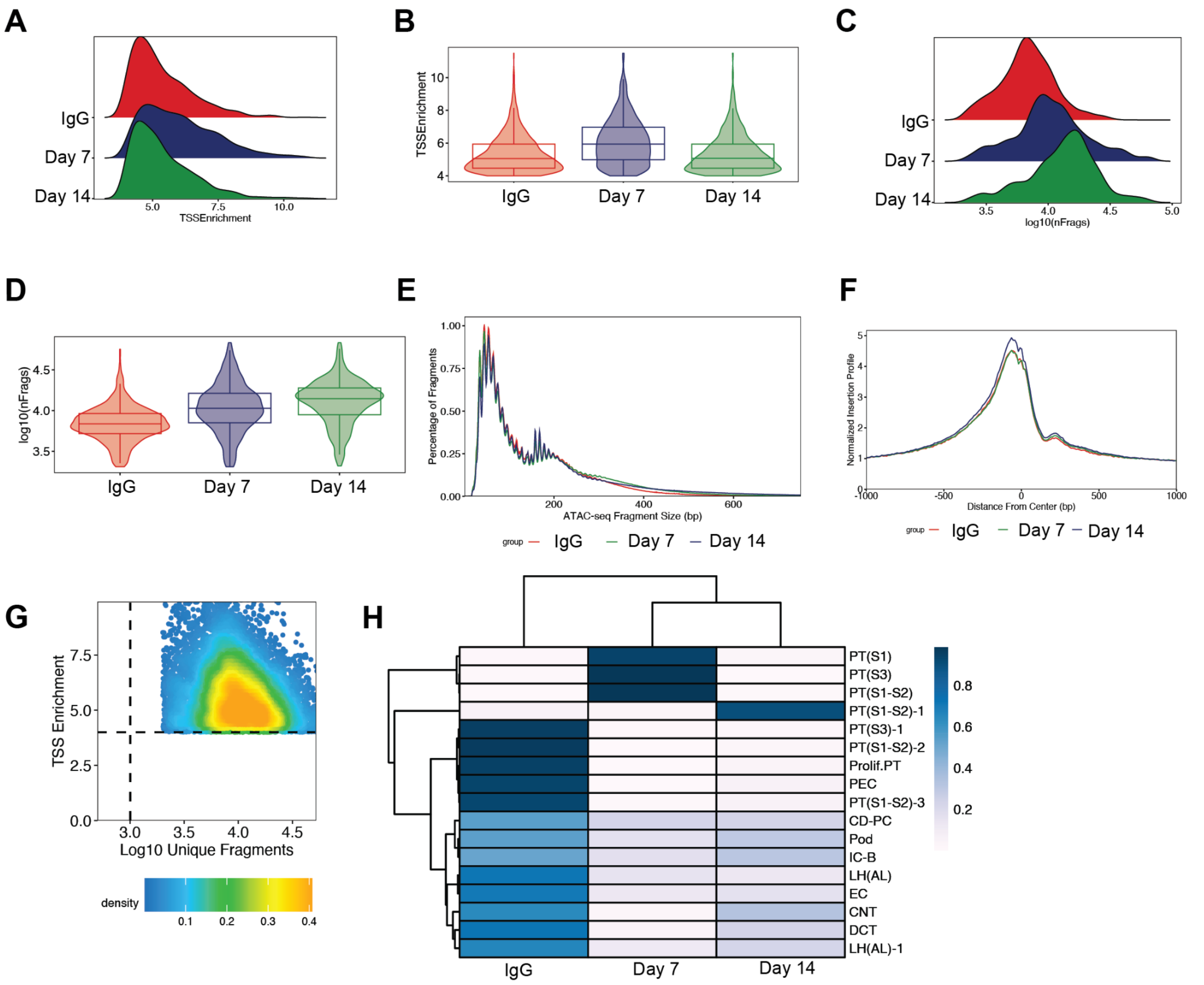


<u>**Supplemental Figure 1**</u>**: 10X Multiome data quality control plots from the NTS model. (A.)** ArchR snATAC-seq TSS enrichment score ridge plots. **(B.)** ArchR snATAC-seq TSS enrichment score violin plots. **(C.)** Ridge plots of unique nuclear fragments. **(D.)** Violin plots of unique nuclear fragments. **(E.)** snATAC-seq fragment size distribution plot. **(F.)** TSS enrichment profile graph centered on the transcription start sites (TSSs) of all genes in the snATAC-seq dataset. **(G.)** Plotting TSS enrichment vs the Log10 unique nuclear fragments displaying cutoffs (dotted lines) for cells used in all downstream analyses. **(H.)** snATAC-seq based confusion matrix heatmap, displaying what proportion of cells occupy which clusters.

### Supplemental Figure 2

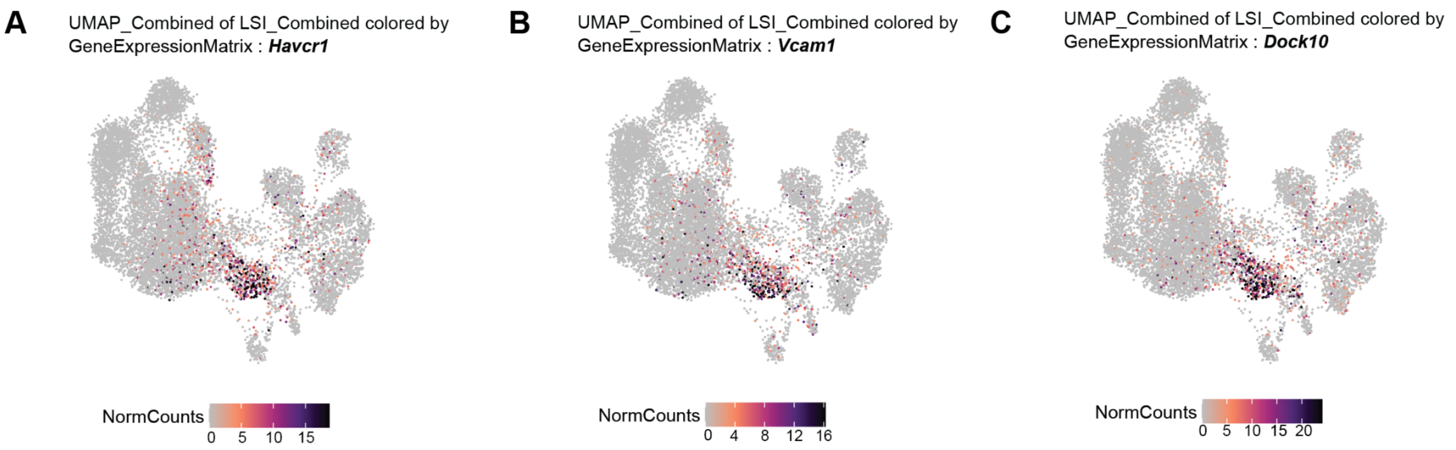


**Supplemental Figure 2: 10X Multiome UMAPs for key PEC-enriched genes in the NTS model.** UMAP for all groups in the NTS model for **(A.)** *Havcr1*, **(B.)** *Vcam1*, and **(C.)** *Dock10* expression.

**Supplemental Figure 3**

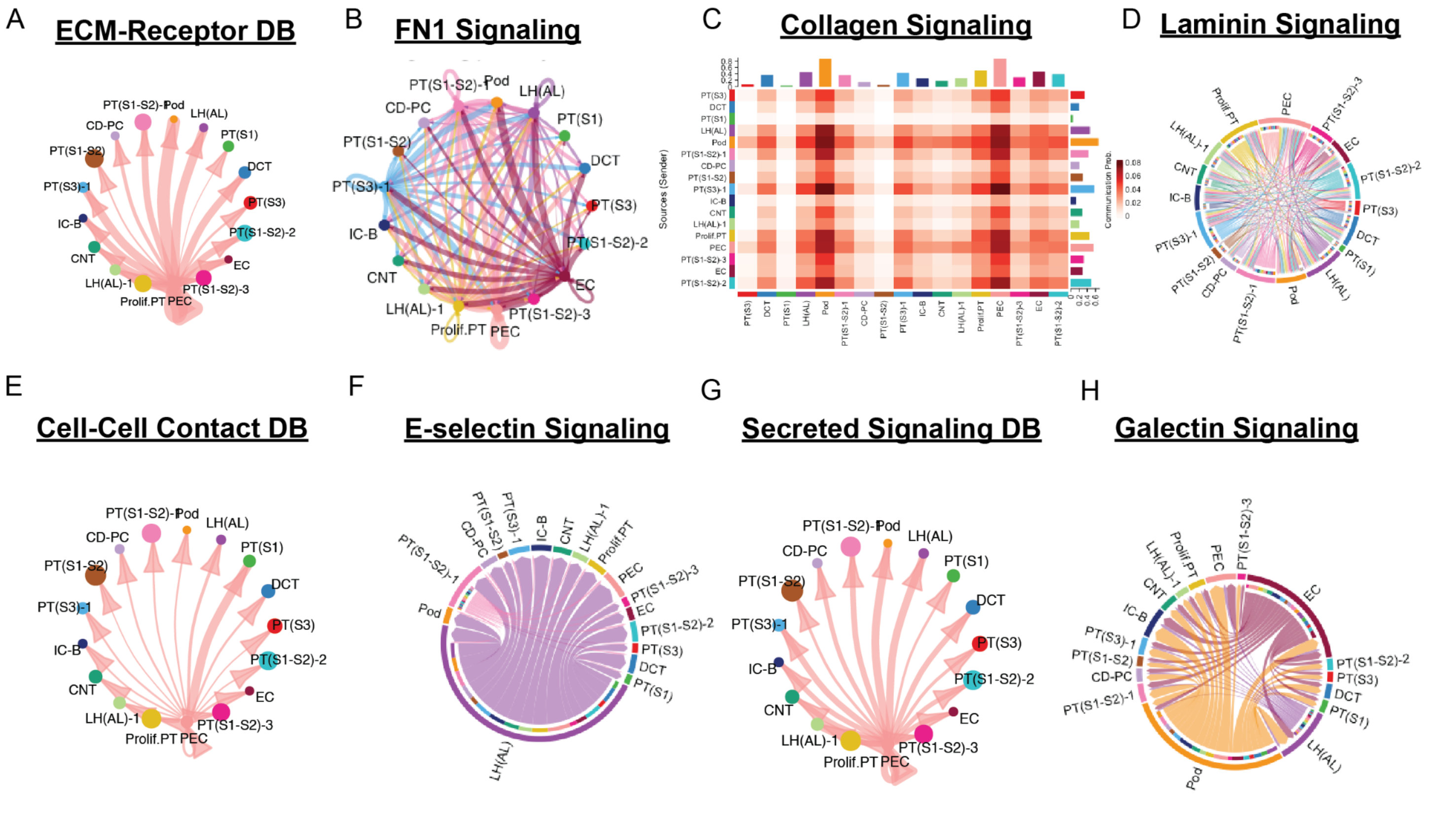


**<u>Supplemental Figure 3</u>: CellChat CD44 ligand/receptor maps from the NTS model. (A.)** CellChat interaction strength (weight) diagram displaying all ECM/Receptor ligand-receptor loops. **(B.)** Cell-cell communication interaction strength diagram of Fibronectin 1 (FN1) signaling pathway network (ECM/Receptor database). **(C.)** COLLAGEN network heatmap of sender/receiver links (ECM/Receptor database). **(D.)** Chord diagram displaying LAMININ signaling pathway network. **(E.)** CellChat interaction strength (weight) diagram displaying all Cell/Cell Contact database derived ligand-receptor loops. **(F.)** Chord diagram displaying E-SELECTIN signaling pathway network (Cell/Cell Contact database). **(G.)** CellChat interaction strength (weight) diagram displaying all Secreted Signaling database derived ligand-receptor loops. **(H.)** CellChat chord diagram displaying GALECTIN signaling pathway network (Secreted Signaling database).

**Supplemental Figure 4**

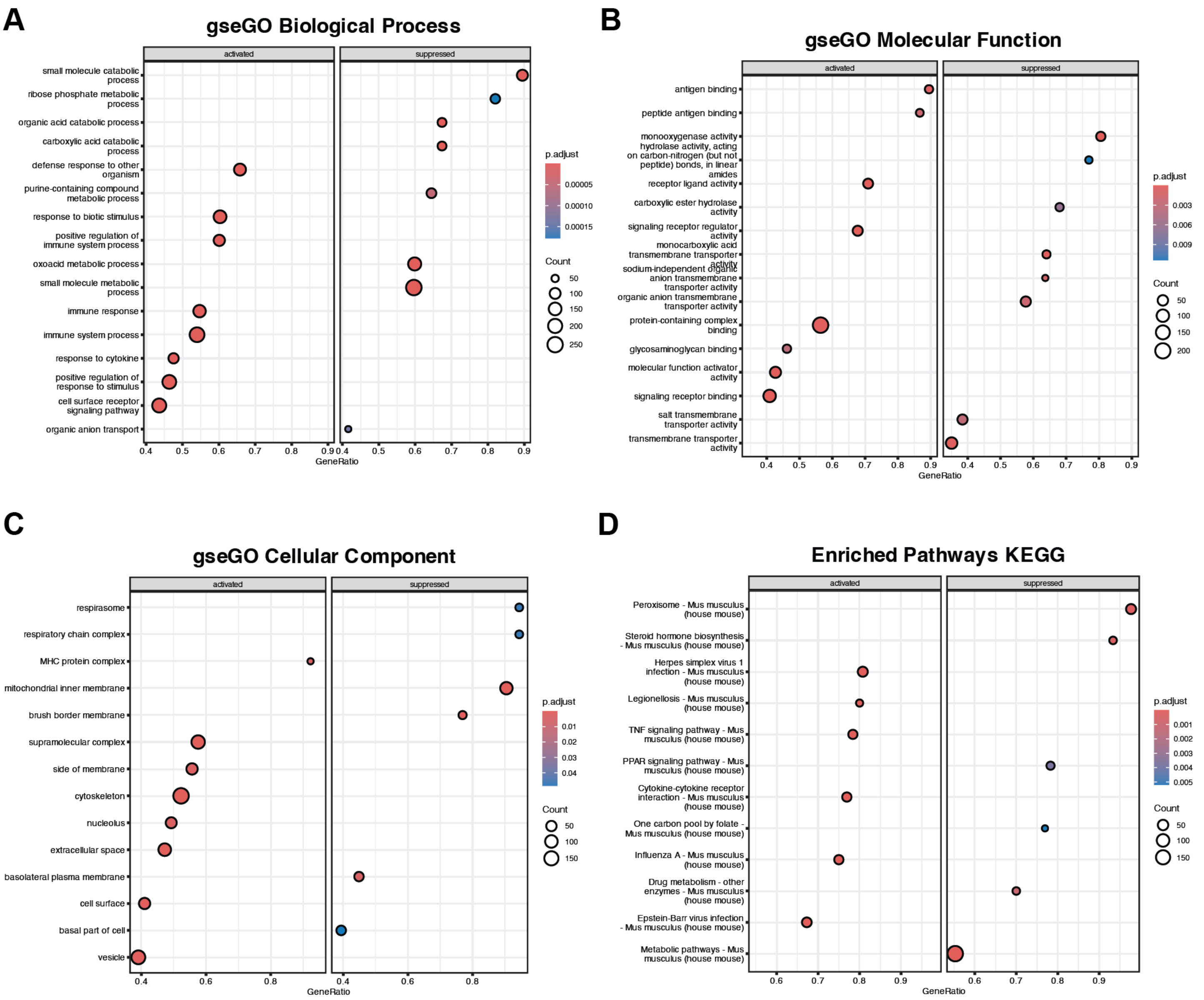


**<u>Supplemental Figure 4</u>: Gene ontology and KEGG pathway GSEA for PEC cluster using the multiome dataset in the NTS model. (A.)** GSEA for GO (gseGO) Biological Process, **(B.)** gseGO Molecular Function using the upregulated and downregulated DEGs, **(C.)** gseGO Cellular Component, and **(D.)** GSEA using KEGG pathway for the upregulated and downregulated DEGs in the PEC cluster.

**Supplemental Figure 5**

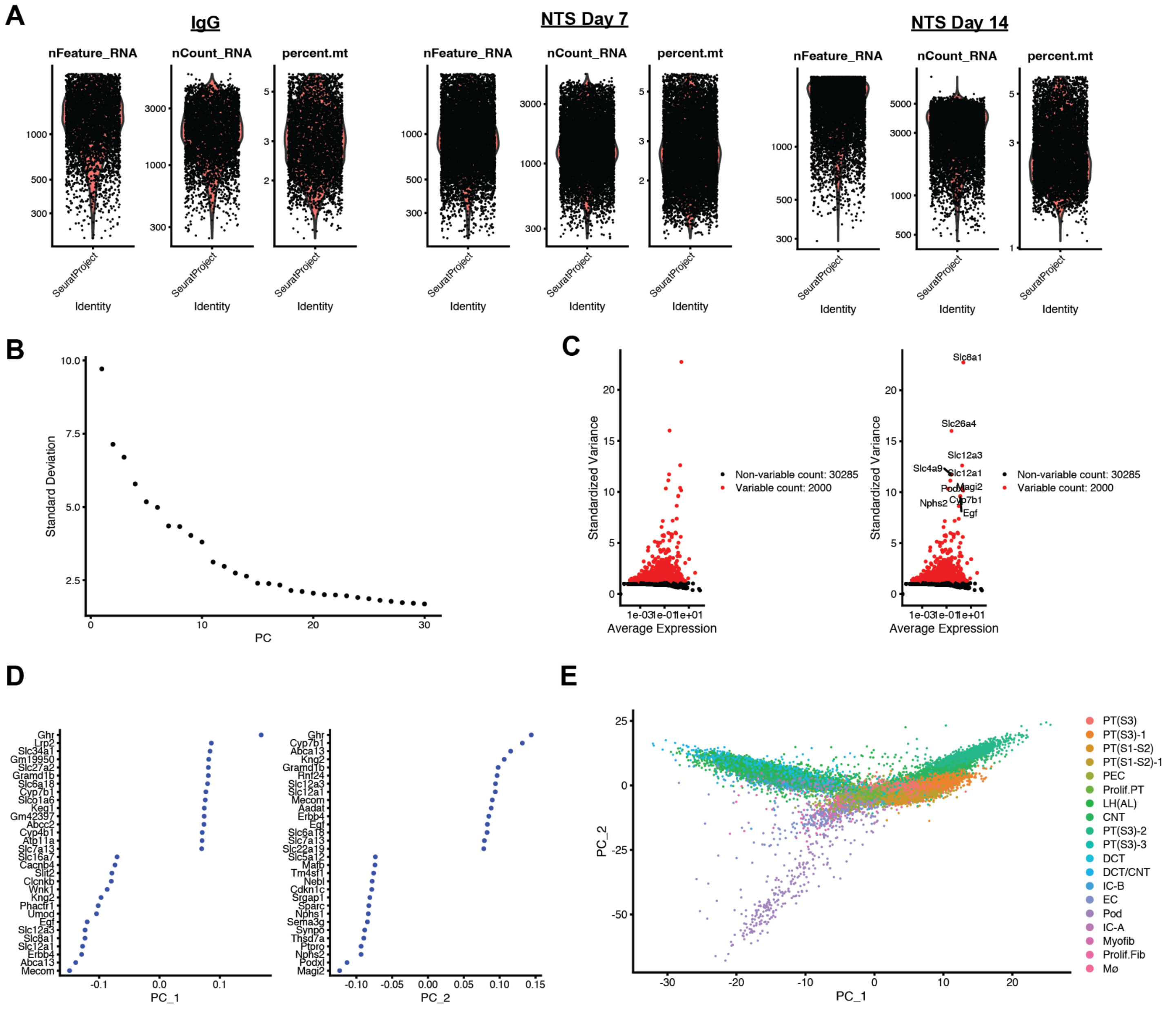


**<u>Supplemental Figure 5</u>: 10X snRNA-seq data quality control plots from the NTS model. (A.)** Seurat generated violin plots of RNA counts/features and percentage of mitochondrially encoded gene transcripts per cell. **(B.)** Elbow plot displaying principal component rankings based on percentage of variance. **(C.)** Scatter plot of highly variable features, plotting the standardized variance against the average expression. **(D.)** Seurat DotPlot displaying gene composition of the first two principal components. **(E.)** UMAP of dimensionality reduction, comparing the first two principal components.

**Supplemental Figure 6**

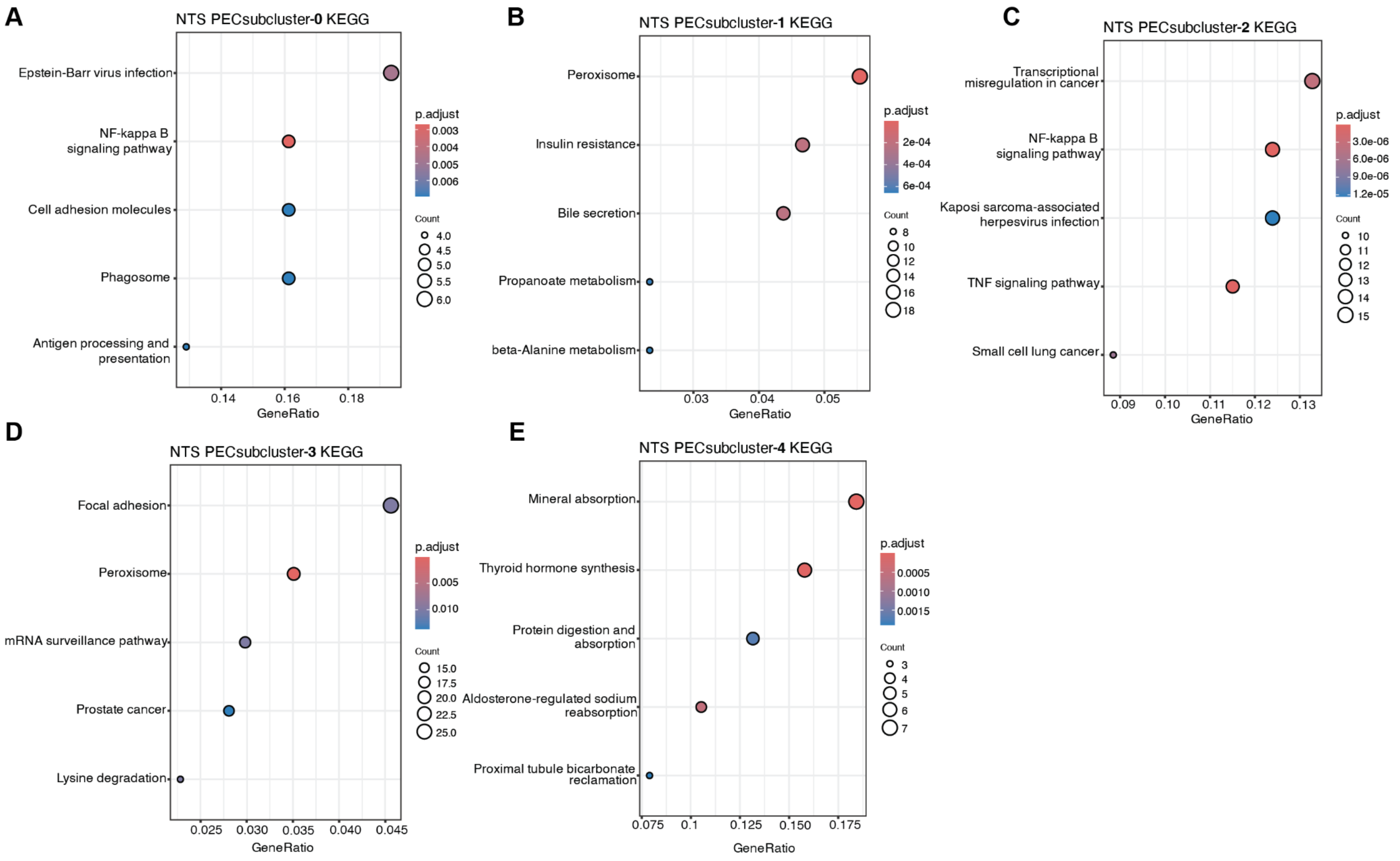


<u>**Supplemental Figure 6**</u>**: KEGG pathway analysis of DEGs from the PEC sub-clusters in the NTS model. (A-E.)** ClusterProfiler generated KEGG pathway analysis DotPlots for each PEC subcluster against all other subclusters.

### Supplemental Figure 7

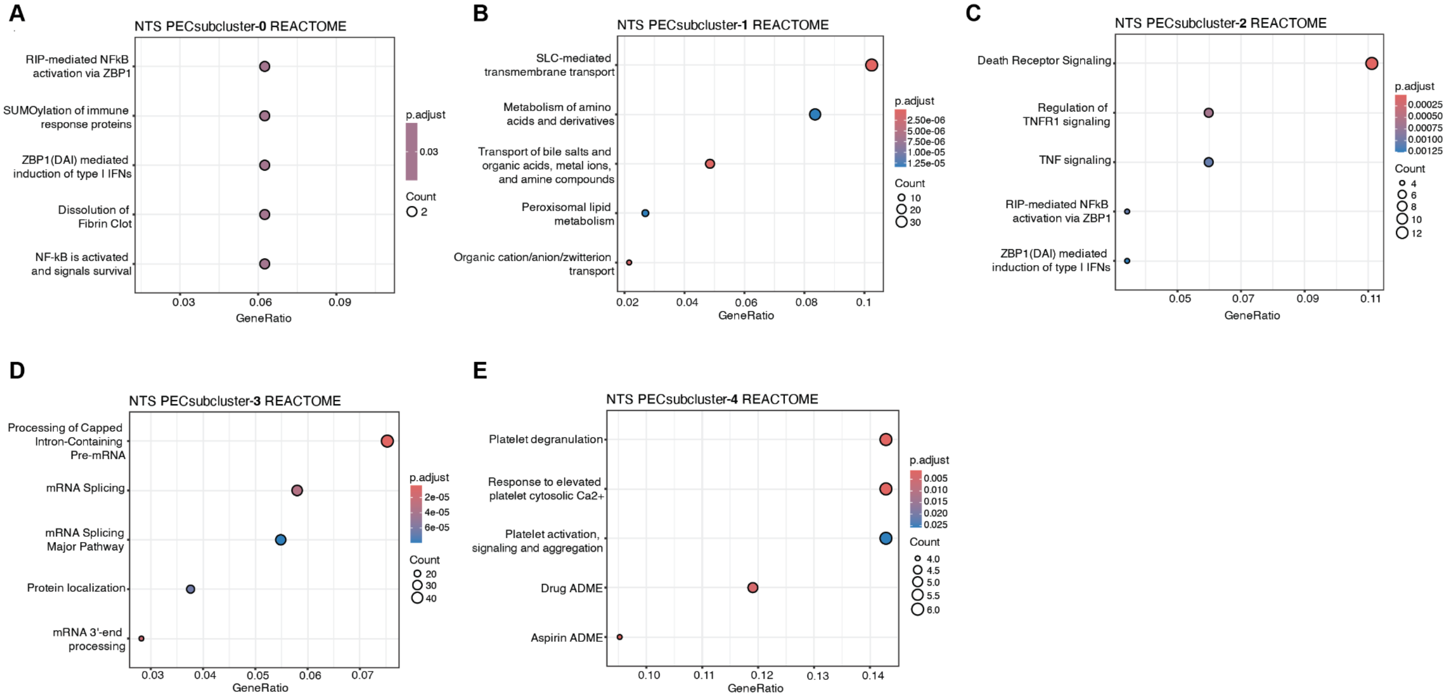


<u>Supplemental Figure 7</u>: **REACTOME pathway analysis of DEGs from the NTS snRNA-seq PEC sub-clusters in the NTS model. (A-E.)** ClusterProfiler generated REACTOME pathway analysis DotPlots for each PEC subcluster against all other subclusters.

**Supplemental Figure 8**

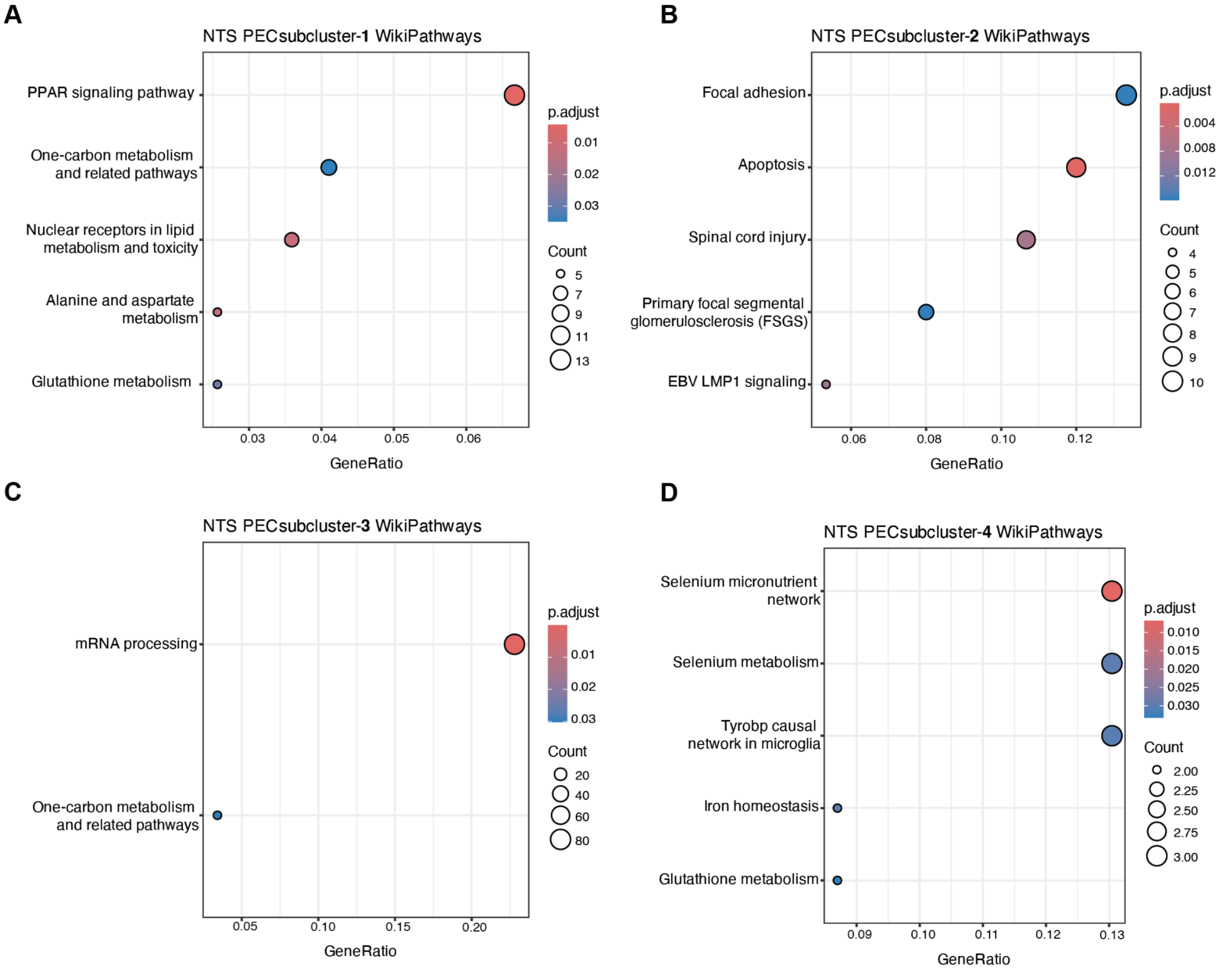


<u>**Supplemental Figure 8**</u>**: WikiPathway analysis of DEGs from the NTS snRNA-seq PEC sub-clusters in the NTS model. (A-D.)** ClusterProfiler generated WikiPathway analysis DotPlots for each PEC subcluster against all other subclusters.

**Supplemental Figure 9**

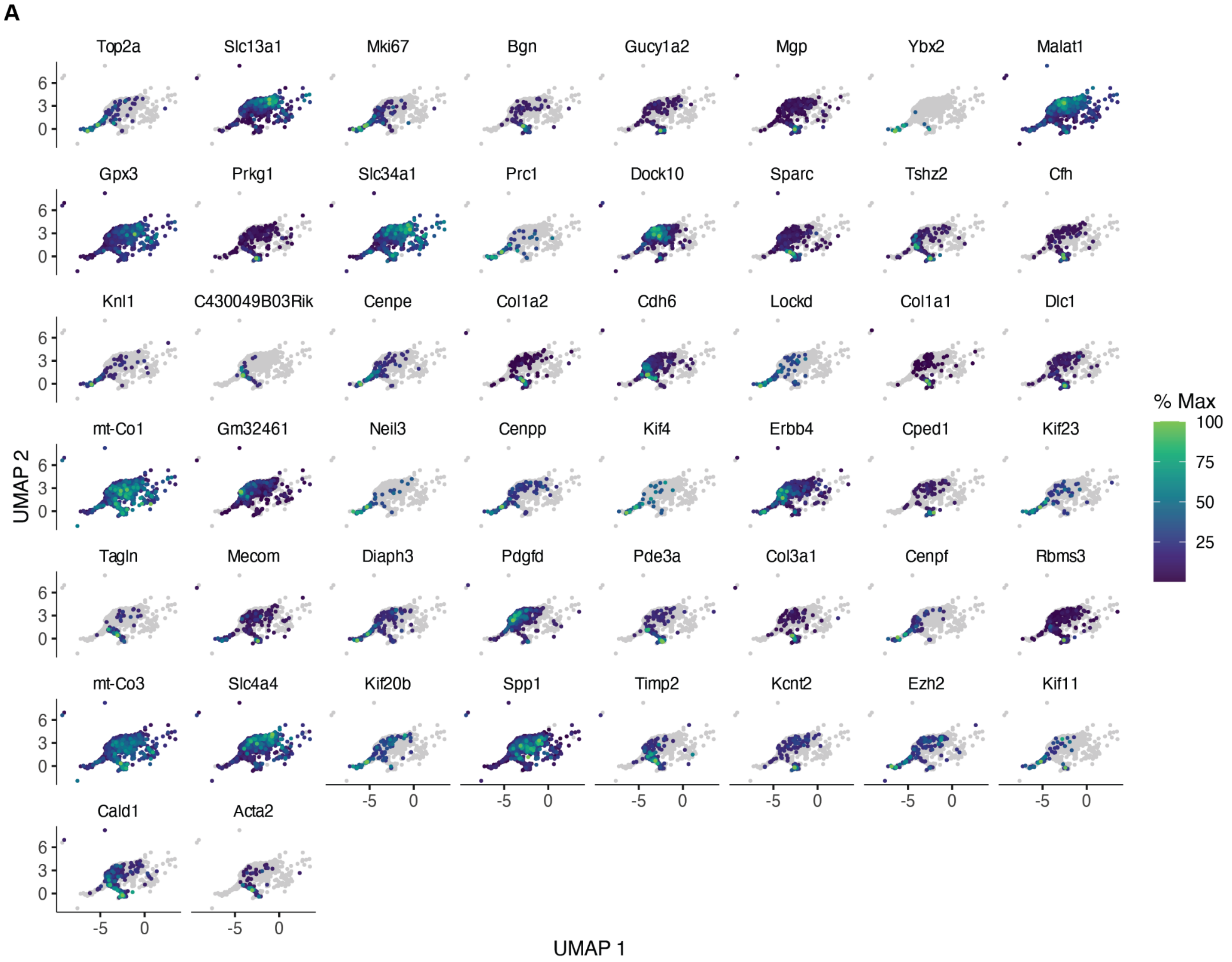


**<u>Supplemental Figure 9</u>: Monocle3 top 50 PEC pseudotime modules in the NTS snRNA-seq dataset.** Top 50 Monocle3 PEC/Prolif. PEC/Myofibroblast module gene marker UMAPs.

### Supplemental Figure 10

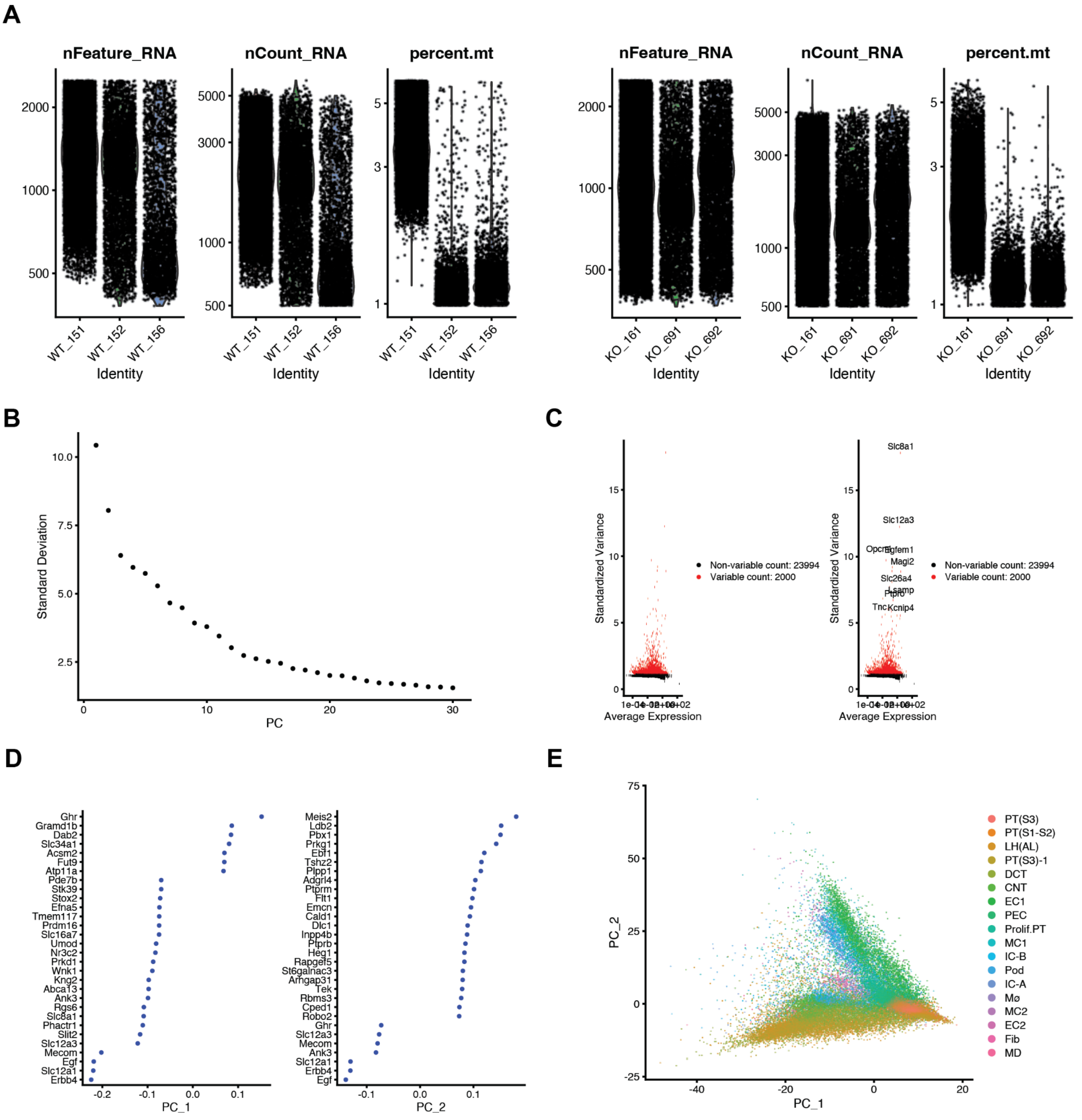

**<u>Supplemental Figure 10</u>: 10X snRNA-seq data quality control plots made in Seurat for *Klf4$^{fl/fl}$* versus *Klf4$^{\Delta Pod}$* mice. (A.)** Seurat generated violin plots of RNA counts/features and percentage of mitochondrially encoded gene transcripts per cell. **(B.)** Elbow plot displaying principal component rankings based on percentage of variance. **(C.)** Scatter plot of highly variable features, plotting the standardized variance against the average expression. **(D.)** Seurat DotPlot displaying gene composition of the first two principal components. **(E.)** UMAP of dimensionality reduction results, comparing the first two principal components.

### Supplemental Figure 11

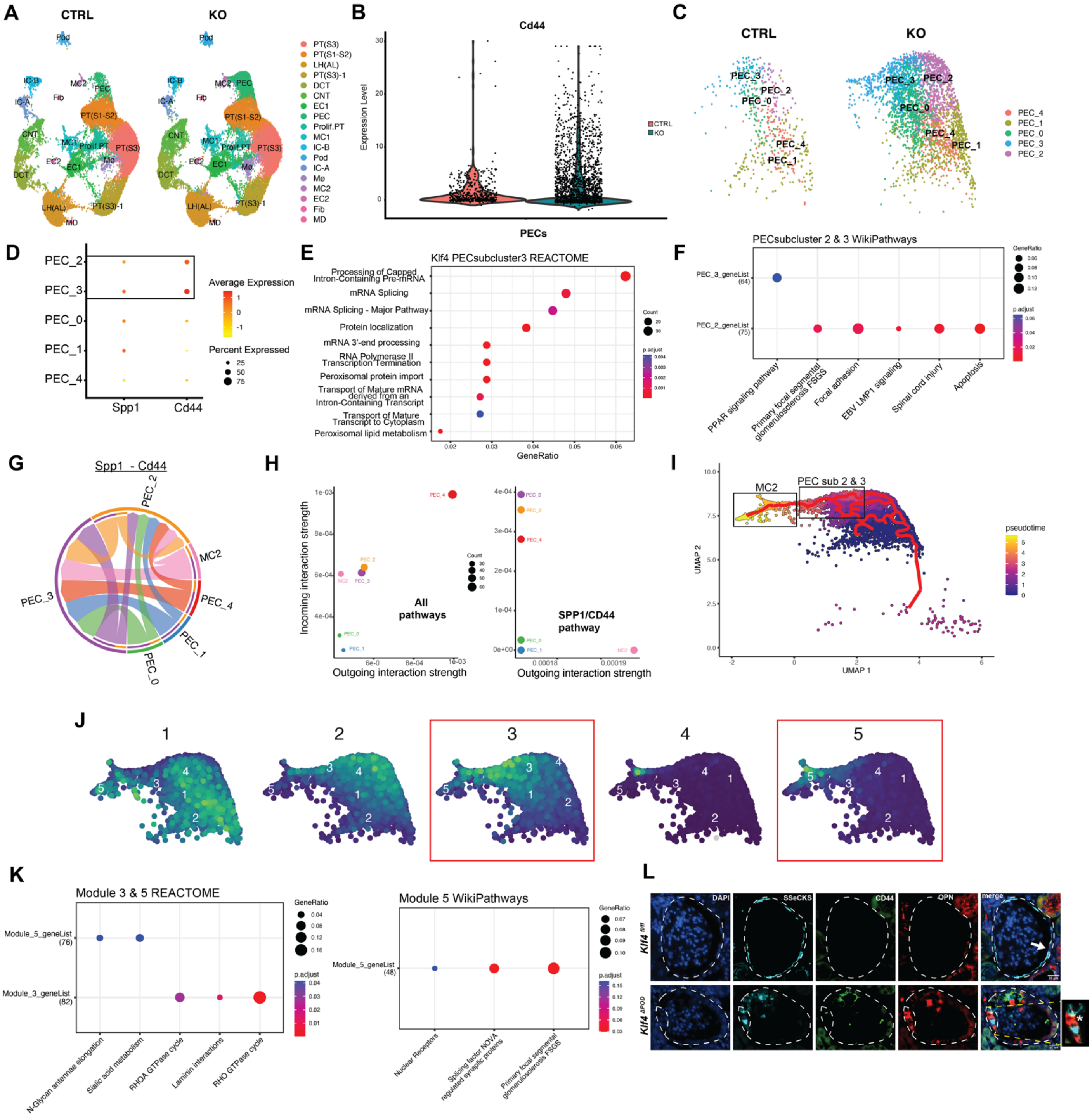

**<u>Supplemental Figure 11</u>: 10X snRNA-seq data analysis (Seurat, CellChat, Monocle3) for *Klf4$^{fl/fl}$* versus *Klf4$^{\Delta Pod}$* mice.** **(A.)** Seurat RNA UMAPs subdivided by both groups. **(B.)** Violin plot for *Cd44* expression in the PEC cluster. **(C.)** UMAP of PEC subclusters. **(D.)** DotPlot of *Cd44* and *Spp1* gene expression across the PEC subclusters. **(E.)** Pathway analysis (*REACTOME*) results for PEC subcluster 3. **(F.)** Pathway analysis (*WikiPathways*) results for PEC subclusters 2 & 3. **(G.)** CellChat CD44-SPP1 ligand receptor interaction analysis utilizing a combination of PEC subclusters and the closely associated mesangial cell 2 (MC2) cluster. **(H.)** CellChat incoming/outgoing interaction strength plots of all possible ligand-receptor pairs (left plot) and CD44-SPP1 specifically (right plot). **(I.)** Monocle3 pseudotime module trajectory across PEC subcluster/MC2 interface, with root cells positioned in PEC subclusters 2 & 3. **(J.)** Top 5 Monocle3 pseudotime trajectory modules determined across PEC subcluster/MC2 interface, with modules 3 & 5 selected based on pseudotime endpoint enrichment in cell populations of interest (PEC subclusters 2 & 3, and MC2). **(K.)** Monocle3 modules 3 & 5 pathway analysis (*REACTOME* & *WikiPathways*). **(L.)** Representative images of immunofluorescence staining for DAPI, SSeCKS, CD44 & OPN in *Klf4$^{fl/fl}$* versus *Klf4$^{\Delta Pod}$* mice (n=3 per group). Dotted white line marks the glomerular region, specifically labeling overlap between PEC-specific markers: 1) <u>*Klf4$^{fl/fl}$*</u>: White arrow marking presence of SSeCKS+ quiescent PECs and, 2) <u>*Klf4$^{\Delta Pod}$*</u>: SSeCKS, CD44 & OPN (white asterisk in merged image pointing to co-localization event) - on the outer perimeter of Bowman's Capsule.

### Supplemental Figure 12

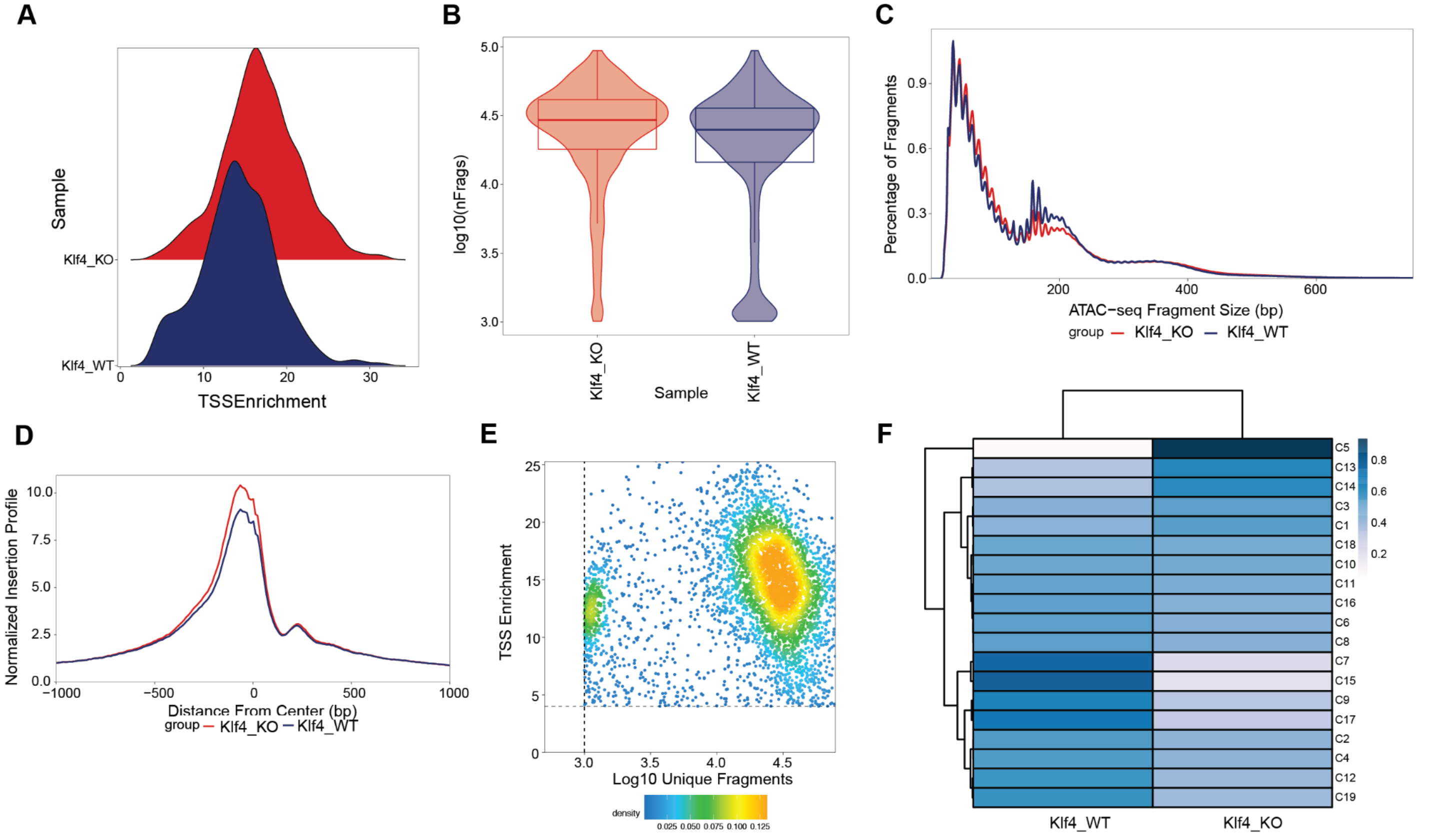


**Supplemental Figure 12: 10X snATAC-seq data quality control plots made in ArchR for *Klf4*$^{fl/fl}$ versus *Klf4*$^{\Delta Pod}$ *mice*. (A.)** Ridge plots of the TSS enrichment score. **(B.)** Violin plots of the unique nuclear fragments (log10(nFrags)). **(C.)** Fragment size distribution plot displaying open chromatin, mono, and di-nucleosomal Tn5 cut regions of the genome. **(D.)** Normalized Tn5 insertion profile +/- 1 kb of the TSS of all genes. **(E.)** Density plot of TSS enrichment scores (y-axis) and unique nuclear fragments (x-axis) displaying per-cell quality distribution. **(F.)** Hierarchical confusion matrix plot across the snATAC-seq dataset, displaying the spread of cluster-specific gene expression across both groups.

**Supplemental Figure 13**

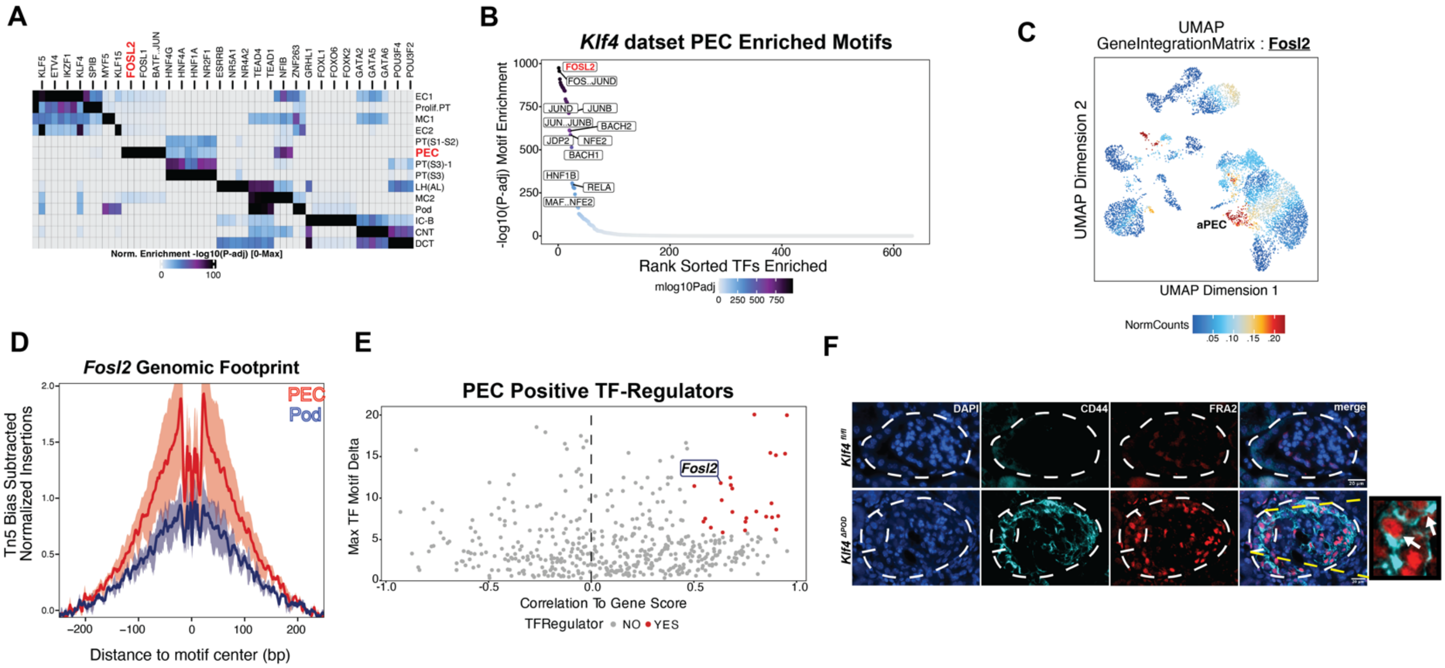


**Supplemental Figure 13: ArchR standard data analysis pipeline with immunofluorescence staining in *Klf4*$^{fl/fl}$ and *Klf4*$^{\Delta Pod}$ mice. (A.)** ArchR heatmap of TF motif enrichment across all clusters. **(B.)** Scatter plot of rank-sorted (-log10(P-adj)) motif enrichment when comparing the ArchR PEC cluster to all other cell clusters. **(C.)** ArchR UMAP feature-plot employing integrated snRNA/ATAC-seq datasets to denote PEC-specific enrichment of *Fosl2* expression. **(D.)** Tn5 bias-normalized snATAC-seq genomic footprinting comparison between podocyte and PEC clusters for *Fosl2* **(E.)** Scatter plot of ArchR called positive TF-regulators (red dots). **(F.)** Representative images of immunofluorescence staining for DAPI, CD44 and FRA2 in *Klf4*$^{fl/fl}$ versus *Klf4*$^{\Delta Pod}$ mice (n=3 per group). Dotted white line marks the glomerular region with the white arrows in enlarged *Klf4*$^{\Delta Pod}$ panel pointing out examples of FRA2/CD44 double-positive activated PECs.

**Supplemental Figure 14**

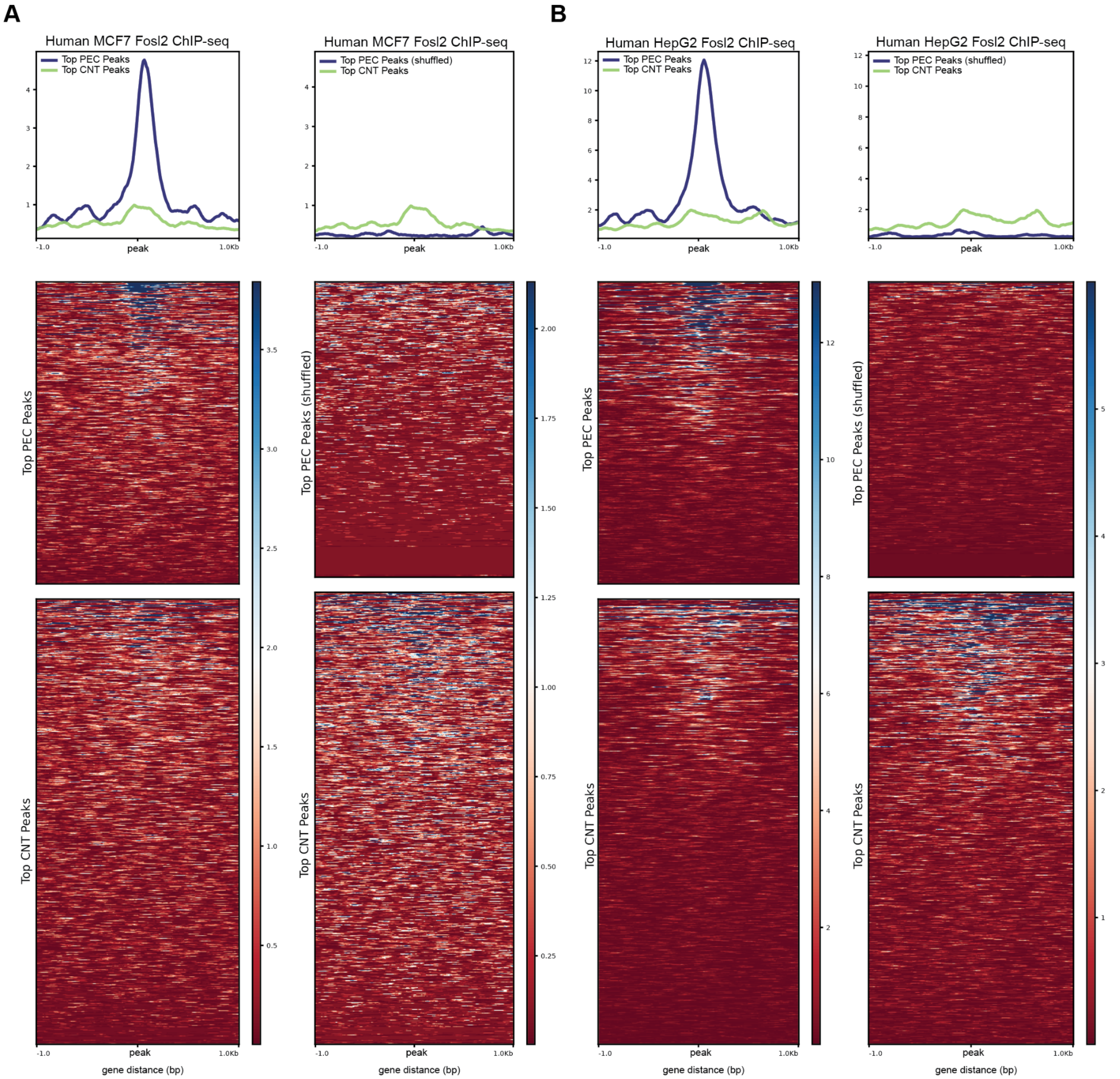


**<u>Supplemental Figure 14</u>: DeepTools profile plots and associated heatmaps of overlap between mouse NTS open chromatin peak locations and FOSL2 ChIP-seq binding sites in human epithelial cell lines. (A.)** Intersect between top mouse PEC and CNT peaks with human MCF7 FOSL2 binding sites (left) and shuffled control PEC peaks (right). **(B.)** Intersect between top mouse PEC and CNT peaks with human HepG2 FOSL2 binding sites (left) and shuffled control PEC peaks (right).
